# A cyclo-DOPA 6-O-glucosyltransferase-mediated route for gomphrenin I biosynthesis in *Basella alba* and *Gomphrena globosa*

**DOI:** 10.64898/2026.04.27.721068

**Authors:** Tomohiro Imamura, Ryouta Shigehisa, Akio Miyazato, Nami Matsumura, Kaisei Miyaki, Tenta Segawa, Masahide Yoshizumi, Hiroki Takagi, Takumi Yamaguchi, Shinya Ohki, Masashi Mori

## Abstract

- Betacyanins are red pigments characteristic of Caryophyllales and show considerable structural diversity, yet the enzymatic basis underlying 6-O-glucosylated betacyanins such as gomphrenin I has remained unclear. In particular, how alternative glucosylation patterns contribute to betacyanin diversification is poorly understood.
- Here, we identified cyclo-DOPA glucosyltransferases from *Basella alba* and *Gomphrena globosa* and examined their roles in gomphrenin I biosynthesis using transient expression assays and tobacco BY-2 cell systems. Phylogenetic analyses, structural modelling and site-directed mutagenesis were employed to investigate their functional and structural characteristics.
- BacDOPA5/6GTs catalysed both 5-O- and 6-O-glucosylation of cyclo-DOPA, leading to the production of betanin and gomphrenin I, whereas GgcDOPA6GT specifically mediated gomphrenin I formation. These enzymes belong to distinct subclades within the cDOPA-GT family, and mutational analyses demonstrated essential roles for conserved histidine residues and an α-helical region adjacent to the catalytic site.
- Thermal stability analyses further showed that gomphrenin I is more thermally stable than betanin, likely due to the formation of an intramolecular hydrogen bond. Together, these results reveal an additional cDOPA6GT-mediated route for gomphrenin I biosynthesis and provide insight into the diversification and functional specialization of betacyanins, linking the position of glucosylation to pigment stability and biochemical properties.

## Introduction

Plant pigmentation plays a vital role not only in visual attraction for pollinators and seed dispersers but also in shielding plants from diverse abiotic stresses, including UV radiation and oxidative damage (Grotewold, 2006; Gould, 2004). Among the various classes of plant pigments, betalains constitute a unique group of water-soluble, nitrogen-containing compounds that are found in plants of the order Caryophyllales (Brockington et al., 2011), where they replace the more widespread anthocyanins in some species, as well as in certain basidiomycete fungi (Eichenberger et al., 2009). Betalains are divided into two major subgroups: betacyanins, which impart red to violet coloration, and betaxanthins, which appear yellow to orange. In contrast to anthocyanins, betalains retain nitrogen atoms within their structures. Notably, betalains and anthocyanins are mutually exclusive in plants, and to date, no species has been shown to synthesize both pigment type simultaneously (Stafford, 1994). Betalains are generally considered to exhibit different stability profiles compared with anthocyanins, particularly under certain pH and temperature conditions (Azeredo, 2009; Gandía-Herrero and García-Carmona, 2013). This physicochemical robustness has been proposed as one possible factor contributing to the prevalence of betalains in certain Caryophyllales species.

Betalains are commonly extracted from plants and used as natural food colorants due to their strong pigmentation. Beyond their application as additives, betalain-containing phytochemicals have been reported to exhibit promising pharmacological potential, particularly in inflammation- and cancer-related contexts (Kapadia et al., 1996; Martinez et al., 2015). To date, about 75 betalains have been identified from plants of about 17 families; in addition, 3 other pigments have been identified from the fly agaric Amanita muscaria (Belhadj Slimen et al., 2017), the biological activities of only a limited subset have been characterized. Among these, betanin (betanidin 5-O-β-glucoside), the major red pigment in beetroot extract, has been shown to induce both apoptosis and autophagic cell death in human cancer cells (Nowacki et al., 2015). Likewise, indicaxanthin, a yellow pigment, exhibits anti-inflammatory activity (Allegra et al., 2014) and exerts antiproliferative and pro-apoptotic effects in human cancer cells (Naselli et al., 2014). In the past, our group explored the physiological activities of several betalains, including HIV-1 protease inhibition by amaranthin (Imamura et al., 2019) and amyloid-β aggregation suppression by betanin and betaxanthins (Imamura et al., 2022, 2025). Consistent with these findings, Martínez-Rodríguez et al. (2024) also reported Aβ aggregation-inhibitory activity for several betaxanthins.

More recently, various betalain biosynthetic pathways have been elucidated (Figures 1a and S1). In general, the betalain biosynthesis pathway begins with hydroxylation of L-tyrosine to form L-3,4-dihydroxyphenylalanine (L-DOPA), a reaction that is often catalyzed by CYP76AD enzymes (Polturak et al., 2016; Sunnadeniya et al., 2016). L-DOPA then acts as a branching substrate, being converted either into betalamic acid by DOPA 4,5-dioxygenase (Christinet et al., 2004; Gandía-Herrero and García-Carmona, 2012) or into cyclo-DOPA by CYP76ADα (Hatlestad et al., 2012). Betalamic acid spontaneously condenses with amino acids to form yellow betaxanthins (Schliemann et al., 1999) or with cyclo-DOPA to generate red-violet betacyanins (Steiner et al., 1999). Further structural diversification of betacyanins proceeds through glucosylation and acylation reactions catalyzed by specific transferases, including those responsible for 5-O-glucosylation of cyclo-DOPA (Sasaki et al., 2005), 5-O- and 6-O-glucosylation of betanidin (Vogt, 2002; Das et al., 2013), the glucuronosyltransferase that synthesizes amaranthin (Imamura et al., 2019), and the recently identified acyltransferase that adds malonic acid to betanin (Glitz et al., 2025).

**Figure 1.**
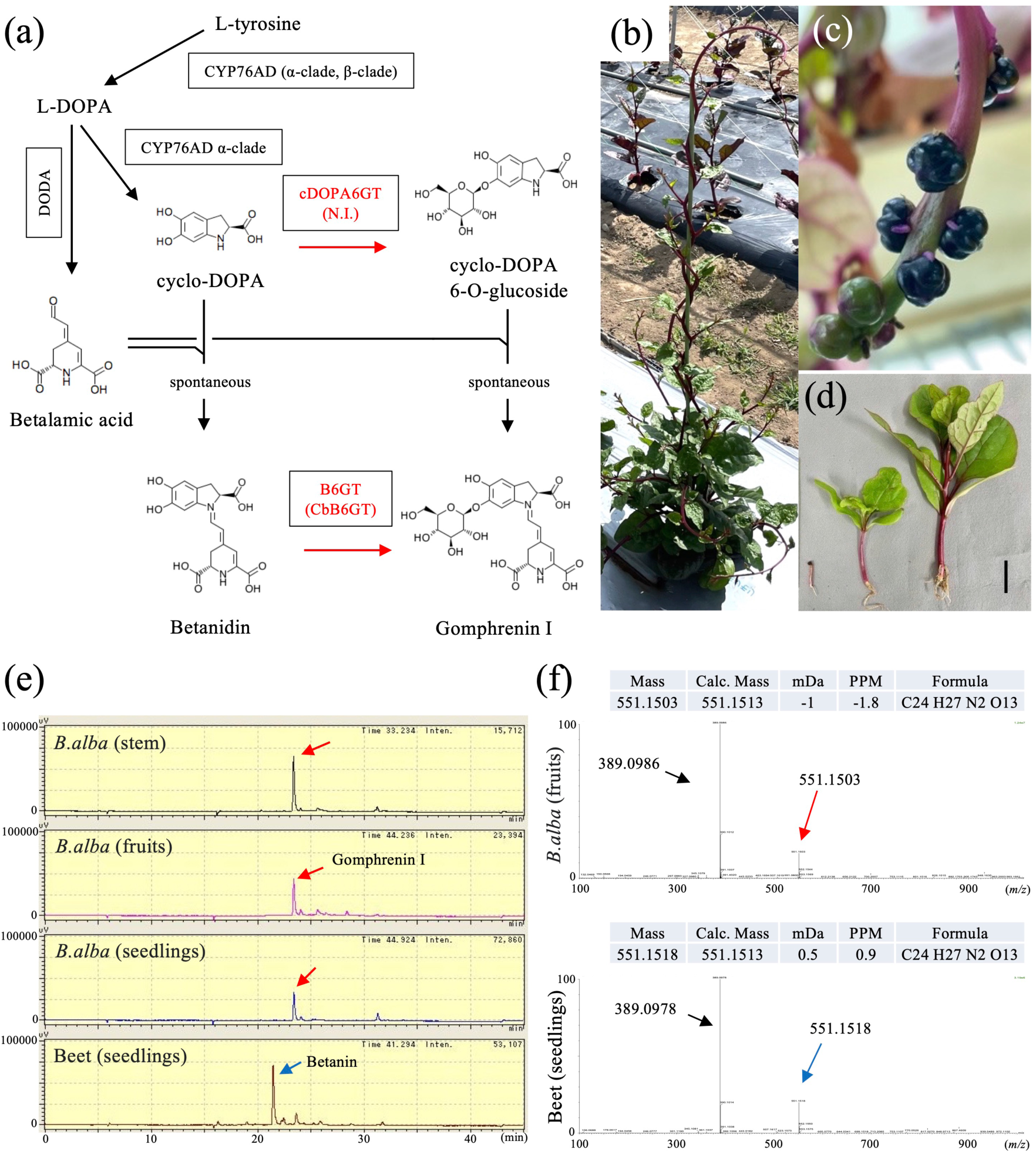
Pigment analysis of *Basella alba* plants. **(a)** Schematic overview of the betacyanin biosynthetic pathway. Boxes indicate enzymes involved in betalain formation, and red arrows highlight the gomphrenin I route. CYP76AD1, cytochrome P450 76AD1; DODA1, DOPA 4,5-dioxygenase 1; cDOPA6GT, cyclo-DOPA 6-O-glucosyltransferase; CbB6GT, betanidin 6-O-glucosyltransferase from *Cleretum bellidiforme*; N.I., not identified. **(b–d)** Photographs of *B. alba* showing plant morphology and betalain accumulation in stems and fruits. **(b)** Field-grown plant. **(c)** Fruits. **(d)** Seedlings (left), a three-week-old plant (middle), and a one-month-old plant (right). Bar, 4 cm. **(e)** HPLC chromatograms of extracts from *B. alba* stems, fruits, and seedlings and from beet seedlings. Red and blue arrows denote gomphrenin I and betanin, respectively. The horizontal axis shows retention time (min), and the vertical axis shows signal intensity (µV). **(f)** Mass spectra corresponding to the HPLC peaks indicated in **(e)**. Upper and lower panels show spectra from *B. alba* fruit extract and beet seedlings, respectively. Red and blue peaks correspond to gomphrenin I and betanin, respectively. The horizontal axis indicates mass-to-charge ratio (*m/z*), and the vertical axis indicates relative abundance. Arrows identify betanidin fragments derived from these pigments.

Betacyanin pigments are classified into several subgroups according to the sugar substitution patterns present on the betanidin aglycone (Kumorkiewicz-Jamro et al., 2021). Of these, gomphrenin-type betacyanins constitute a distinct subgroup characterized by the core structure of gomphrenin I, in which a β-D-glucose moiety is O-glycosidically attached to the 6-hydroxyl position of betanidin (Kumorkiewicz-Jamro et al., 2021). Representative plant species that predominantly accumulate gomphrenin-type pigments include *Basella alba* and *Gomphrena globosa* (Minale et al., 1967; Lin et al., 2010). In this biosynthetic route, the enzyme that produces the core compound gomphrenin I is betanidin 6-O-glucosyltransferase (B6GT), which catalyzes the transfer of a glucose moiety to the 6-hydroxyl group of betanidin (Figure S2). The corresponding *B6GT* gene was originally isolated from cultured cells of *Cleretum bellidiforme* (formerly *Dorotheanthus bellidiformis*), commonly known as Livingstone daisy (Vogt et al., 2002) (Figure 1a). Although these cultured cells are capable of synthesizing gomphrenin I, its accumulation remains low, with betanin persisting as the predominant betacyanin (Heuer et al., 1996). As a result, genes encoding the enzymes responsible for gomphrenin I biosynthesis have not previously been identified in plant species such as *B. alba* and *G. globosa*, in which gomphrenin-type betacyanins constitute the major pigments.

In this study, we focus on *B. alba* and *G. globosa*, two plant species known to produce gomphrenin-type betacyanins. *B. alba* (Malabar spinach) is a climbing plant native to tropical Asia that is widely cultivated across tropical and subtropical regions. *B. alba* accumulates red pigments in various tissues, including fruits, stems, and leaves (Figure 1b–d). *G. globosa* (globe amaranth) is native to Central and South America and is grown worldwide as an ornamental plant. *G. globosa* primarily accumulates red pigments in the bracts. Despite the ornamental and phytochemical significance of these species, genomic information for *B. alba* and *G. globosa* remains unavailable.

To clarify the biosynthetic route leading to gomphrenin-type betacyanins, we set out to identify the genes required for gomphrenin I formation. Through the isolation and functional characterization of *cDOPA5GT* orthologs from *B. alba* and *G. globosa*, we identified enzymes capable of catalyzing 6-O-glucosylation of cyclo-DOPA. These results reveal functional diversity within the cDOPA glucosyltransferase family and provide a framework for the future development of heterologous platforms for the controlled production of gomphrenin-type pigments.

## Materials and Methods

### Plant materials and growth conditions

Seeds of the purple stem variety *of B. alba* was obtained from Sakata Seed Co., Japan, and seed of the *G. globosa* cultivars Audray Purple Red was obtained from Takii Seed Co., Japan. *B. alba* and *G. globosa* seeds were sown in a cell tray and then grown in a phytotron at 23°C under a 12-h light/12-h dark photoperiod. Tobacco BY-2 cells were maintained at 26°C in Linsmaier and Skoog medium supplemented with 3% sucrose and 0.2 mg/L 2,4-dichlorophenoxyacetic acid (Nagata *et al*., 1992).

### RNA-seq analysis

RNA-seq analysis was performed to obtain comprehensive mRNA sequence and expression profiles in *B. alba* red leaves and *G. globosa* involucral bracts. Total RNA was extracted from gomphrenin I-accumulating tissues using the RNeasy Plant Mini Kit (Qiagen, Valencia, CA, USA) and treated with RNase-free DNase I (Qiagen) to remove genomic DNA contamination. For Illumina sequencing, 1 µg of total RNA was used for library preparation following the manufacturer’s protocol for the NEBNext Poly(A) mRNA Magnetic Isolation Module (NEB, Ipswich, MA, USA). Libraries were subjected to 150 bp paired-end sequencing on a NovaSeq 6000 platform (Illumina, San Diego, CA, USA). Raw sequence reads in FASTQ format were filtered for quality prior to analysis. Gene expression levels were compared with RNA-seq datasets from other studies by converting read counts to transcripts per kilobase million values (Wagner et al., 2012).

### Molecular cloning

Total RNA was extracted using the RNeasy Plant Mini Kit (Qiagen) and then treated with RNase-free DNase I (Qiagen) to eliminate residual genomic DNA. First-strand cDNA was synthesized from 500 ng of total RNA using the PrimeScript II 1st Strand cDNA Synthesis Kit (TaKaRa, Kusatsu, Japan) with oligo(dT) primers. Full-length open reading frame (ORF) sequences of *BaB6GTL1*, *BaB6GTL2*, *BacDOPA5/6GT1*, *BacDOPA5/6GT2*, and *GgcDOPA6GT* were obtained from the RNA-seq dataset (Table S1). Next, the full-length ORFs of *CbB6GT*, *PucDOPA5GT*, and *SmcDOPA5/6GT* were synthesized as codon-optimized genes for tobacco expression (Genscript Biotech, Nanjing, China). Site-directed mutagenesis was performed using a PCR-based method with mutation-specific primers (Table S3), followed by nested PCR to construct the desired mutant constructs.

### RT-PCR analysis

First-strand cDNA was synthesized from 500 ng of total RNA using the PrimeScript II 1st Strand cDNA Synthesis Kit (TaKaRa) with oligo(dT) primers. RT-PCR was performed on GeneAtlas 322 (Astec, Fukuoka, Japan) using PrimeSTAR GXL DNA Polymerase (TaKaRa). Amplification of the candidate transcripts consisted of an initial denaturation at 94°C for 2 min followed by 35 cycles at 98°C for 10 s, 55°C for 15 s, and 68°C for 1.5 min. *L23* and *NtCesA* served as positive controls for the expression in *N. benthamiana* leaves and tobacco BY-2 cells, respectively (Imamura *et al*., 2019). Primer pairs are listed in Table S3.

### Plasmid construction

PrimeSTAR GXL DNA polymerase and oligonucleotides containing the appropriate restriction enzyme cleavage sites were used for PCR amplification (Table S3). For agro-infiltration analysis in *N. benthamiana*, the amplified fragments of the candidate genes and the corresponding mutation constructs involved in gomphrenin I biosynthesis were digested with relevant restriction enzymes and then introduced into the binary vector pCAMBIA1301MdNcoI (Imamura *et al*., 2018). Additional expression vectors (i.e., pCAM-CYP76AD1-1, pCAM-CqDODA-1, pCAM-CqcDOPA5GT, and pCAM- AcGFP1) had been generated previously (Imamura *et al*., 2018). For the stable transformant analysis using BY-2 cells, the amplified fragments of *CqCYP76AD1-1* and *CqDODA-1* were digested with the appropriate restriction enzymes and introduced into the binary vector pBI121 (Imamura *et al*., 2019). The resulting plasmids were then sequenced using BigDye terminator chemistry and an ABI PRISM 3,100 genetic analyzer (Applied Biosystems, Foster City, CA, USA).

### Transient expression in *Nicotiana benthamiana*

Expression constructs were introduced into *Agrobacterium tumefaciens* strain GV3101 using the triparental mating method (Wise *et al*., 2006). The resulting *Agrobacterium* suspensions were then infiltrated into leaves of 5- to 6-week-old *Nicotiana benthamiana* plants following the procedure described previously (Shamloul *et al*., 2014). After infiltration, the plants were maintained in a growth chamber at 23°C and 60% humidity under long-day conditions (16-h light/8-h dark).

### Phylogenetic tree of deduced amino acid sequences

We used ClustalW to align deduced amino acid sequences of cyclo-DOPA 5-O-glucosyltransferase homologs from a variety of plant species (Thompson *et al*., 1994) (Table S4). A phylogenetic tree was then constructed using the neighbor-joining algorithm implemented in MEGA11 (Tamura *et al*., 2021).

### Structural analyses

Three-dimensional structures of the proteins were modeled using the AlphaFold2 web server (Jumper et al., 2021). The crystal structure of the *Trollius chinensis* C-glycosyltransferase (TcCGT1; PDB ID: 6JTD) was used as a reference structure because it represents one of the few plant family-1 glycosyltransferases with a resolved crystal structure and shares the conserved fold and PSPG motif characteristic of plant UGTs. The reference structure was used to guide structural interpretation and to facilitate comparison of the active site architecture among plant UGTs. Overall, 97% of amino acid residues were included in the modeled region.

### Transformation of BY-2 cells

Tobacco BY-2 cells were grown in Linsmaier and Skoog medium supplemented with 3% sucrose and 0.2 mg L⁻¹ 2,4-dichlorophenoxyacetic acid at 26°C (Nagata et al., 1992). *Agrobacterium* tumefaciens strain GV3101 harboring the respective binary vectors was used for transformation following a previously described procedure (Hagiwara et al., 2003). Transgenic calli were selected on agar medium containing the appropriate selective agents, namely 50 mg L⁻¹ hygromycin, and 100 mg L⁻¹ kanamycin, together with 500 mg L⁻¹ carbenicillin. For each construct, more than ten independent transgenic BY-2 cell lines were generated. From these, two independent lines showing comparable and stable pigment accumulation were selected for detailed analysis. Suspension cultures derived from selected calli were initially grown in 3 mL liquid medium in six-well plates for primary screening and were subsequently transferred to 150 mL liquid medium in 500-mL flasks with constant shaking at 135 rpm. After an initial culture period of 2–3 weeks, the suspension cells were maintained without selective agents. Cells were examined using an Axiovert 200 optical microscope (Zeiss, Jena, Germany). All images were captured using Axiovision version 4.6 (Zeiss).

### Plant pigment chemical analysis

Pigments were extracted from *N. benthamiana* leaves and BY-2 cells using the method described previously (Imamura *et al*., 2019). Briefly, tissues were homogenized in an aqueous solvent at room temperature. The crude extracts were then mixed with an equal volume of acetonitrile and centrifuged to remove precipitated impurities. The clarified supernatants were then concentrated using a centrifugal concentrator (CC-105, Tomy Seiko Inc., Tokyo, Japan) to remove acetonitrile and concentrate the pigment extracts. A Shimadzu LC-20AD system (Kyoto, Japan) was used for analytical HPLC separations. Samples were then loaded onto a Shim-pack GWS C18 column (5 µm; 200 × 4.6 mm i.d.; Shimadzu GLC, Tokyo, Japan), and linear gradients were applied from 0% B to 45% B over 45 min using 0.05% trifluoroacetic acid (TFA) in water (solvent A) and 0.05% TFA in acetonitrile (solvent B). The flow rate was 0.5 mL min^−1^ at 25°C and elution was monitored by absorbance at 536 nm.

Next, infected leaves were collected and pigments were extracted to quantify betacyanin production in *N. benthamiana*. To do so, extracts were first dried using a centrifugal concentrator and subsequently dissolved in ultrapure water at a ratio of 30 µL per 10 mg fresh leaf weight. After dissolution, the samples were analyzed by HPLC, and peak areas corresponding to the target betacyanin compounds were measured. Relative production levels were calculated by setting the peak area of the wild-type sample to 100%. All values represent the mean of three biological replicates (N = 3). Error bars indicate the mean ± SD. *p* < 0.05 vs. WT (Student’s t-test).

### LC-MS analysis

A Shimadzu LC-20AD system equipped with an electrospray ionization Fourier transform ion cyclotron resonance mass spectrometer (Solarix, Bruker Daltonics) operating in the positive mode was used to perform LC-MS analysis. An XBridge C18 column (150 × 2.1 mm, 3.5-µm particle size; Waters) was used for chromatographic separation. The flow rate was kept at 0.3 mL/min using 0.1% TFA in acetonitrile. A stepwise gradient was applied with 0%, 10%, 50%, and 100% acetonitrile at 0–3, 3–15, 15–20, and 20–25 min, respectively.

### UV−Vis spectroscopy

A UV-2450 spectrophotometer (Shimadzu) was used to perform UV-Vis measurements. Betalain pigment concentration was determined using molar extinction coefficients of ε = 65,000 M^−1^ cm^−1^ at 536 nm for betacyanins (Gandía-Herrero *et al*., 2010). All measurements were carried out in water at 25°C.

### Betacyanins purification

Next, we purified betacyanins (gomphrenin I and betanin) produced in tobacco BY-2 cells for subsequent NMR analysis. Betacyanins were first extracted from BY-2 cells using water as the extraction solvent. The aqueous extract was purified by anion-exchange chromatography (DEAE Sepharose Fast Flow, Cytiva, Uppsala, Sweden), followed by reversed-phase chromatography (COSMOSIL 75C18-OPN, Nacalai Tesque, Kyoto, Japan) according to the method described by Henarejos-Escudero et al. This eluate was evaporated to dryness, and the resulting residue was dissolved in water. This concentrated betacyanin solution was then subjected to separation using a HPLC system (Gilson, Middleton, WI, USA). Samples were then loaded onto a COSMOSIL 5C18 AR-II column (5 µm; 250 × 20 mm i.d.; Nacalai Tesque). A linear gradient from 0% to 50% solvent B was applied over 50 min using 0.05% TFA in water (solvent A) and 0.05% TFA in acetonitrile (solvent B) at a flow rate of 5 mL min⁻¹. Elution was monitored at 536 nm, and we collected fractions corresponding to the elution peaks of betanin and gomphrenin I. These eluates were then evaporated to dryness, after which the resulting residues were dissolved in water and stored at −20°C until further use.

### NMR-based structural analysis of the red pigments

Three-dimensional structures of the red pigments were analyzed by using a Bruker 500 MHz NMR spectrometer equipped with an AVANCE III console and a BBO probe. The sample temperature was maintained at 26°C during NMR experiments. For resonance assignment, one-dimensional ^1^H and ^13^C spectra and several two-dimensional datasets, including DQF-COSY, NOESY, HMBC, and ^1^H-^13^C HSQC, were recorded. DSS was used as the chemical shift reference. Finally, deuterated methanol containing a small amount of deuterated TFA served as the solvent for the NMR samples.

### Quantitative assessment of thermal stability by UV–Vis spectroscopy

To evaluate the thermal stability of betanin and gomphrenin I, approximately 230-250 µM solutions of each pigment were prepared. Heat treatments were carried out for 3 h using a thermal cycler (GeneAtlas G, Astec, Tokyo, Japan) equipped with a gradient function at the following temperatures: 27.7, 28.8, 31.2, 35.2, 40.1, 45.0, 49.9, 54.8, 59.7, 63.7, 66.1, and 67.2°C. Pigment concentrations after each heat treatment were determined by UV-Vis spectroscopy. The experiment was repeated three times. Statistical differences between samples treated at the same temperature were evaluated by Student’s *t*-test. Asterisks denote significant differences between group means (*p* < 0.05).

Structural analysis of thermal stability by ^1^H-NMR spectroscopy Next, we examined the thermal stability of the red pigments via ^1^H-NMR. During this analysis, the sample temperature was gradually increased in 5°C increments from room temperature to 72°C. A waiting time of approximately 15 min was introduced at each temperature to allow thermal equilibration before data acquisition.

### Computational chemistry

MD simulations were performed in explicit solvent at 300 K using the AMBER software package with the GAFF2 force field. The molecule was solvated in a TIP3P water box, and simulations were run for 600 ns under the *NPT* ensemble. Resulting trajectories were analyzed for intramolecular hydrogen bonds using the cpptraj module. A representative structure obtained from this analysis was used as the initial model for subsequent DFT calculations. DFT computations were then performed using Gaussian 16 program at the B3LYP/6-311+G(d,p) level of theory with the Polarizable Continuum Model and water as the solvent.

## Results

### Pigment analysis of *Basella alba*

To identify the gene responsible for gomphrenin I biosynthesis, betacyanins were extracted from *B. alba* and analyzed by high-performance liquid chromatography (HPLC). This analysis confirmed that *B. alba* produces gomphrenin I in its stems, fruits, and seedlings (Figure 1e and f). The retention times of gomphrenin I and betanin were approximately 23.5 min and 21.5 min, respectively, under the HPLC conditions used. Beet extracts, which contain only betanin, were used as a control to validate peak assignment. The resulting chromatographic profiles clearly separated gomphrenin I from betanin (Figure 1e). The identity of the gomphrenin I peak was confirmed based on a combination of evidence, including comparison with beet extracts (which contain only betanin), and LC–MS analysis showing a molecular ion consistent with gomphrenin I (m/z ≈ 551). These results are also consistent with previous reports describing gomphrenin I as the major betacyanin in *B. alba* (Lin et al., 2010). This assignment is further supported by NMR-based structural identification of gomphrenin I in the BY-2 system.

### Identification of candidate genes involved in gomphrenin I biosynthesis in *Basella alba*

A betanidin 6-O-glucosyltransferase (B6GT) involved in gomphrenin I biosynthesis has previously been isolated from *C. bellidiforme* (Vogt et al., 2002). However, alternative biosynthetic routes and the genes responsible for gomphrenin I formation in other species remain unclear. To identify genes involved in gomphrenin I biosynthesis, RNA-seq analysis was performed using *B. alba* red leaves that accumulated gomphrenin I, and transcriptome data were generated from these red-pigmented samples. Using these sequences, a BLASTx search was conducted with the amino acid sequence of the betanidin 6-O-glucosyltransferase from *C. bellidiforme* (CbB6GT; NCBI Accession No. AF374004) as the query. In this study, we identified two candidate genes, *BaB6GTL1* and *BaB6GTL2* (Table S1). Although multiple UGT-like sequences were identified in the transcriptome, candidates were prioritized based on sequence similarity to known B6GT and their expression levels in betalain-producing tissues (Table S1). Relative to CbB6GT, the amino acid sequence identities of BaB6GTL1 and BaB6GTL2 with CbB6GT were 63% and 62%, respectively. Moreover, the two *B. alba* genes shared 94% identity with one another (Figure S3). Next, we evaluated the enzymatic activities of BaB6GTL1 and BaB6GTL2. Our group previously produced the betacyanins betanin and amaranthin in *N. benthamiana* leaves by transient coexpression of quinoa betalain biosynthetic genes (i.e., *CqCYP76AD1-1*, *CqDODA-1*, *CqCDOPA5GT*, and *CqAmaSy*) (Imamura et al., 2018; Imamura et al., 2019). *CqAmaSy,* an amaranthin synthase from *Chenopodium quinoa*, was used as a reference glycosyltransferase. Using this established transient expression system, we tested whether the two *B. alba* candidate genes possessed B6GT activity capable of generating gomphrenin I. Expression plasmids were constructed for each candidate gene as well as for CbB6GT and CqB5GT (NCBI Accession No. XM_021897654), and introduced into *Agrobacterium tumefaciens* (Figure S4). The resulting strains, together with *CqCYP76AD1-1* and *CqDODA-1*, were co-infiltrated into *N. benthamiana* leaves. No red pigmentation was observed in leaves infiltrated with either *BaB6GTL1* or *BaB6GTL2* (Figure S5). An additional control experiment showed that UGT-free infiltration (*CqCYP76AD1-1* + *CqDODA-1* + *P19*) resulted in only very weak pigmentation and trace-level peaks, comparable to those observed in the presence of AcGFP1 (Figure S6; see also Figure S5 for comparison). In contrast, coexpression of *CqB5GT* produced red pigmentation (betanin), whereas *CbB6GT* did not induce visible pigment accumulation (Figure S5). Transgene expression in the infiltrated tissues was confirmed at the transcript level by RT-PCR (Figure S5; corresponding uncropped gel images are provided in Figure S17). These results indicate that expression of *BaB6GTL1* and *BaB6GTL2* did not result in detectable gomphrenin I accumulation under the conditions tested, suggesting that an alternative biosynthetic route may be involved.

Two distinct biosynthetic routes have been proposed for betanin formation (Figure S1), involving either glucosylation of betanidin by B5GT or glucosylation of cyclo-DOPA by cDOPA5GT. Based on this framework, we hypothesized that gomphrenin I biosynthesis may proceed via a similar route mediated by cyclo-DOPA 6-O-glucosyltransferase (cDOPA6GT) (Figure 1a) and attempted to isolate the corresponding gene from *B. alba*. Using the *B. alba* RNA-seq dataset, we performed a BLASTx search using the amino acid sequence of *CqcDOPA5GT* from quinoa (*C. quinoa*, NCBI Accession No. XM_021892614) as a query, a cDOPA5GT known to generate cyclo-DOPA 5-O-glucoside. Two candidate genes, *BacDOPA5/6GT1* and *BacDOPA5/6GT2*, were identified (Table S1). Although multiple UGT-like sequences were identified in the transcriptome, candidates were prioritized based on sequence similarity to previously characterized cDOPA glucosyltransferases and their expression levels in betalain-producing tissues (Table S1). Both sequences shared 61% amino acid identity with *CqcDOPA5GT*, and the two *B. alba* genes exhibited 92% identity with each other (Figure S7).

Next, we assessed the activities of *BacDOPA5/6GT1* and *BacDOPA5/6GT2* using a transient expression assay in *N. benthamiana* leaves. Expression plasmids for each gene (Figure S4) were introduced into *A. tumefaciens*, and the resulting cultures were co-infiltrated with *CqCYP76AD1-1* and *CqDODA-1* into *N. benthamiana* leaves. Red pigmentation developed in leaves infiltrated with either *BacDOPA5/6GT1* or *BacDOPA5/6GT2* (Figure 2a). Transgene expression in the infiltrated tissues was confirmed by RT-PCR (Figure 2b). Pigments extracted from the red-colored leaves were analyzed by HPLC. Both the BacDOPA5/6GT1 and BacDOPA5/6GT2 proteins produced compounds with retention times identical to those of plant-derived betanin and gomphrenin I (Figure 2c). Taken together, these findings demonstrate that BacDOPA5/6GT1 and BacDOPA5/6GT2 possess both cDOPA5GT activity for betanin biosynthesis and cDOPA6GT activity for gomphrenin I biosynthesis. BacDOPA5/6GT2 exhibited lower activity than BacDOPA5/6GT1 under the same experimental conditions, although both enzymes produced betanin and gomphrenin I.

**Figure 2.**
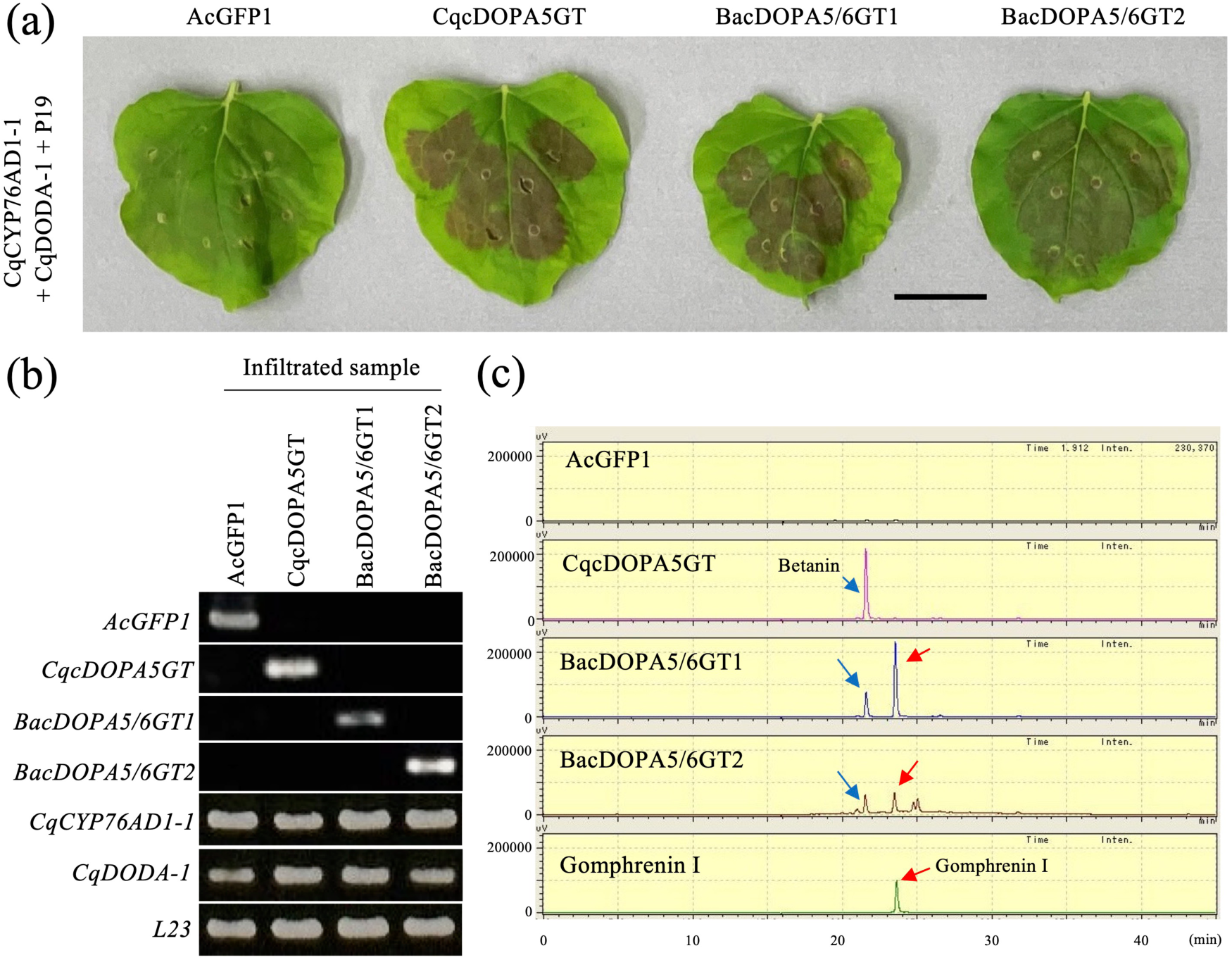
Functional characterization of *BacDOPA5/6GT* from *Basella alba*. **(a)** Recombinant expression of *BacDOPA5/6GTs* and *CqcDOPA5GT* in *N. benthamiana* leaves. *Agrobacterium* strains carrying BacDOPA5/6GT1, BacDOPA5/6GT2, or CqcDOPA5GT were co-infiltrated with CqCYP76AD1- 1, CqDODA-1, and P19. AcGFP1 served as the negative control. Bar, 4 cm. **(b)** RT-PCR analysis of transgene expression in infiltrated leaves, with *L23* serving as the internal control. **(c)** HPLC chromatogram of pigments extracted from infiltrated *N. benthamiana* leaves. Blue and red arrows indicate the peaks corresponding to betanin and gomphrenin I, respectively. The horizontal axis shows retention time (min), and the vertical axis shows signal intensity (µV).

### BacDOPA5/6GTs contribute to gomphrenin I biosynthesis

Since our results strongly suggest that BacDOPA5/6GT1 and BacDOPA5/6GT2 may possess cDOPA6GT activity and that they may be linked to gomphrenin I production, we determined the molecular structure of their products by nuclear magnetic resonance (NMR) spectroscopy. Because the structure of betanin has been established previously, NMR analysis in this study was focused on confirming the identity of gomphrenin I produced in the heterologous system. Moreover, since NMR analysis requires a relatively large amount of pigment, we used a previously established BY-2 protocol (Imamura et al., 2019). Using this system, we constructed BY-2 cell lines expressing *BacDOPA5/6GT* to facilitate structural identification of the putative gomphrenin I compound.

Plasmids carrying different drug resistance markers were constructed to overexpress the *CqCYP76AD1*, *CqDODA-1*, and *BacDOPA5/6GT* genes (Figure S8). The resulting *Agrobacterium tumefaciens* transformants were prepared and used to introduce these constructs into BY-2 cells. Three stable cell lines were generated: a betanidin-producing line harboring *CqCYP76AD1-1* and *CqDODA-1*; a *BacDOPA5/6GT1*-expressing line containing *CqCYP76AD1-1*, *CqDODA-1*, and *BacDOPA5/6GT1*; and a *BacDOPA5/6GT2*-expressing line harboring *CqCYP76AD1-1*, *CqDODA-1*, and *BacDOPA5/6GT2* (Figures 3a and S8b). After confirming transgene expression by RT-PCR (Figure 3b), a strongly pigmented line was selected and maintained in liquid culture. The *BacDOPA5/6GT*-expressing lines consistently displayed vivid red coloration, whereas the betanidin-producing line remained orange and failed to develop red pigmentation throughout the study (Figure 3a). Similar pigment accumulation profiles were observed across additional independent lines (Figure S7).

**Figure 3.**
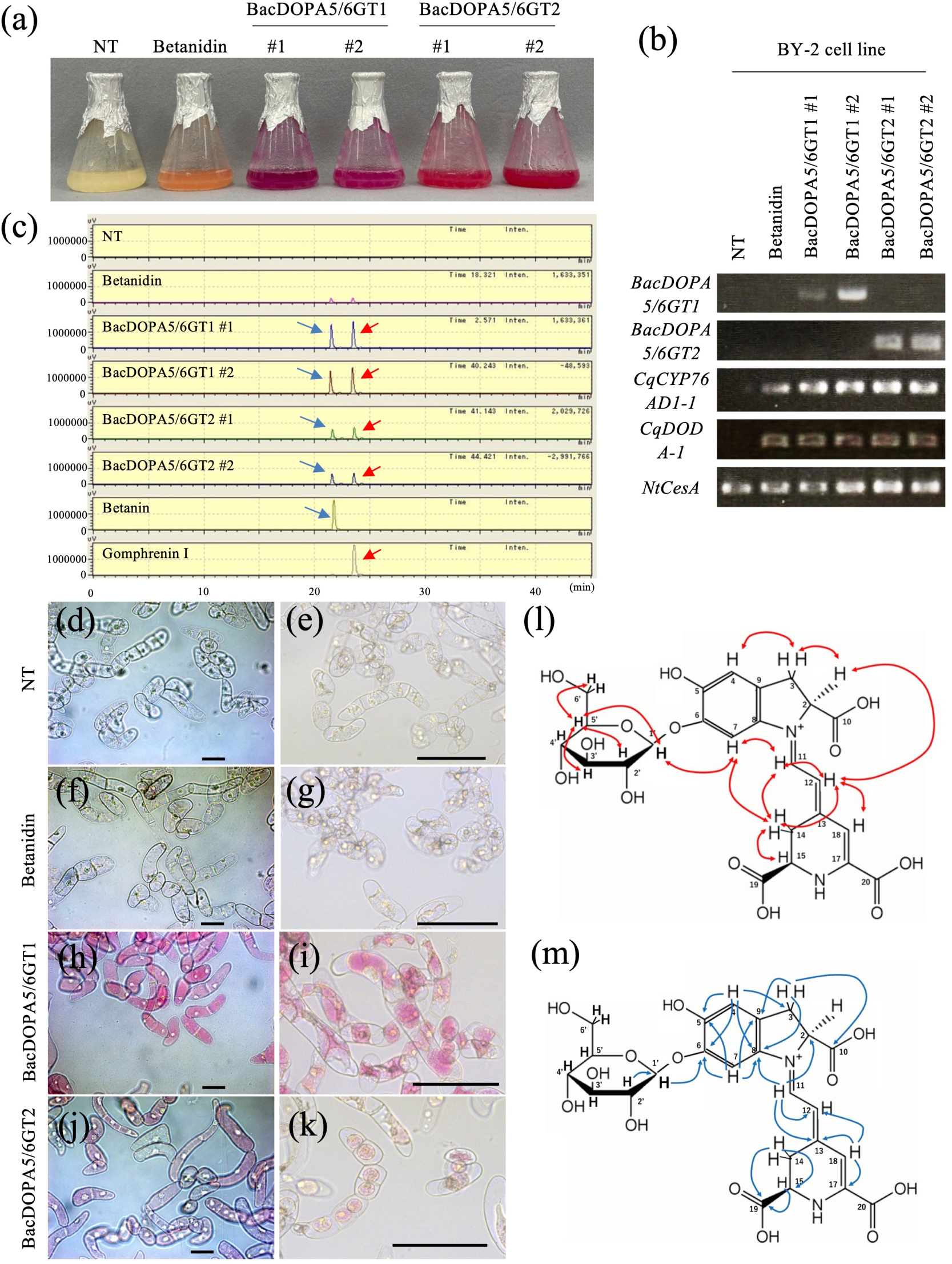
Gomphrenin I production in tobacco BY-2 cell lines. **(a)** Photographs of transformed tobacco BY-2 cell lines two weeks following subculturing. BacDOPA5/6GT1 and BacDOPA5/6GT2 indicate lines expressing each gene together with CqCYP76AD1-1 and CqDODA-1. “Betanidin” denotes line expressing only CqCYP76AD1-1 and CqDODA-1. “NT” indicates the non-transgenic BY-2 cell line. #1 and #2 represent independent transformants. **(b)** RT-PCR analysis of transgene expression, with *NtCesA* serving as the internal control. **(c)** HPLC chromatograms of pigments from transformed lines, with blue and red arrows indicate betanin and gomphrenin I, respectively. The horizontal axis shows retention time (min), and the vertical axis shows signal intensity (µV). **(d-k)** Microscopy images of BY-2 cells: NT cells **(d, e)**, betanidin-producing line **(f, g)**, BacDOPA5/6GT1 #1 line **(h, i)**, and BacDOPA5/6GT2 #1 line **(j, k)**. Panels **(d, f, h, j)** show actively growing cells, whereas panels **(e, g, i, k)** show BY-2 cells after hypertonic treatment. Bars, 100 µm. **(l, m)** Schematic representation of NMR correlations detected in NOESY **(l)** and HMBC **(m)** spectra of gomphrenin I produced by the BacDOPA5/6GT1 BY-2 cell line. Red and blue arrows mark cross-peaks observed in NOESY and HMBC, respectively.

HPLC analysis revealed that the *BacDOPA5/6GT*-expressing lines contained peaks corresponding to betanin and a compound with the same retention time as gomphrenin I, mirroring the pattern observed in the *N. benthamiana* transient expression assay (Figure 3c). Microscopic observation showed that betacyanins accumulated within the vacuoles of all red-colored cell lines (Figure 3d–k). The peak corresponding to the putative gomphrenin I was collected from the *BacDOPA5/6GT1*-expressing BY-2 line and purified. Structural elucidation by NMR spectroscopy confirmed that this compound was gomphrenin I (Figures 3l and m, Figure S9 and Table S5). This result aligned with the previous NMR characterization of gomphrenin I extracted from *B. alba* reported by Wu et al. (2013). These findings indicate that *BacDOPA5/6GTs* possess both cDOPA5GT and cDOPA6GT activities, catalyzing gomphrenin I biosynthesis via cDOPA 6-O-glucoside. The *BacDOPA5/6GT1*-expressing BY-2 line produced 27.1 ± 7.21 µM of betanin and 34.2 ± 10.3 µM of gomphrenin I, whereas the *BacDOPA5/6GT2*-expressing line produced 12.1 ± 0.49 µM of betanin and 12.5 ± 0.9 µM of gomphrenin I. BacDOPA5/6GT2 exhibited lower activity than BacDOPA5/6GT1 under the same experimental conditions, consistent with observations from the *N. benthamiana* transient expression assay. Minor peaks corresponding to betanin and gomphrenin I were occasionally detected in control samples (betanidin-producing line in BY-2 cells), likely due to endogenous glycosyltransferase activities in the host systems, as supported by HPLC and MS analysis (Figure S10). Importantly, LC–MS analysis confirmed that these peaks correspond to glycosylated betacyanins, whereas free betanidin was not detected (Figure S10), indicating that betanidin does not accumulate as a stable intermediate *in vivo*. Consistent with this, LC–MS analysis of the betanidin-producing BY-2 cell line independently showed that the detected pigments correspond to glycosylated derivatives (betanin and gomphrenin I).

### Functional characterization of GgcDOPA6GT from *Gomphrena globosa*

To identify genes involved in gomphrenin I biosynthesis in *G. globosa*, we first analyzed the flower bracts by HPLC and MS, which confirmed that this species produces gomphrenin I (Figure 4). We then carried out RNA-seq analysis using these flower bracts to obtain transcriptome data. From this dataset, a BLASTx search was performed with the amino acid sequence of *CqcDOPA5GT* as the query. One candidate gene, *GgcDOPA6GT*, was identified (Table S1), and its amino acid sequence shared 68% identity with *CqcDOPA5GT* (Figure S7). Although multiple UGT-like sequences were identified in the transcriptome, candidates were prioritized based on sequence similarity to previously characterized cDOPA glucosyltransferases and their expression levels in betalain-producing tissues (Table S1).

**Figure 4.**
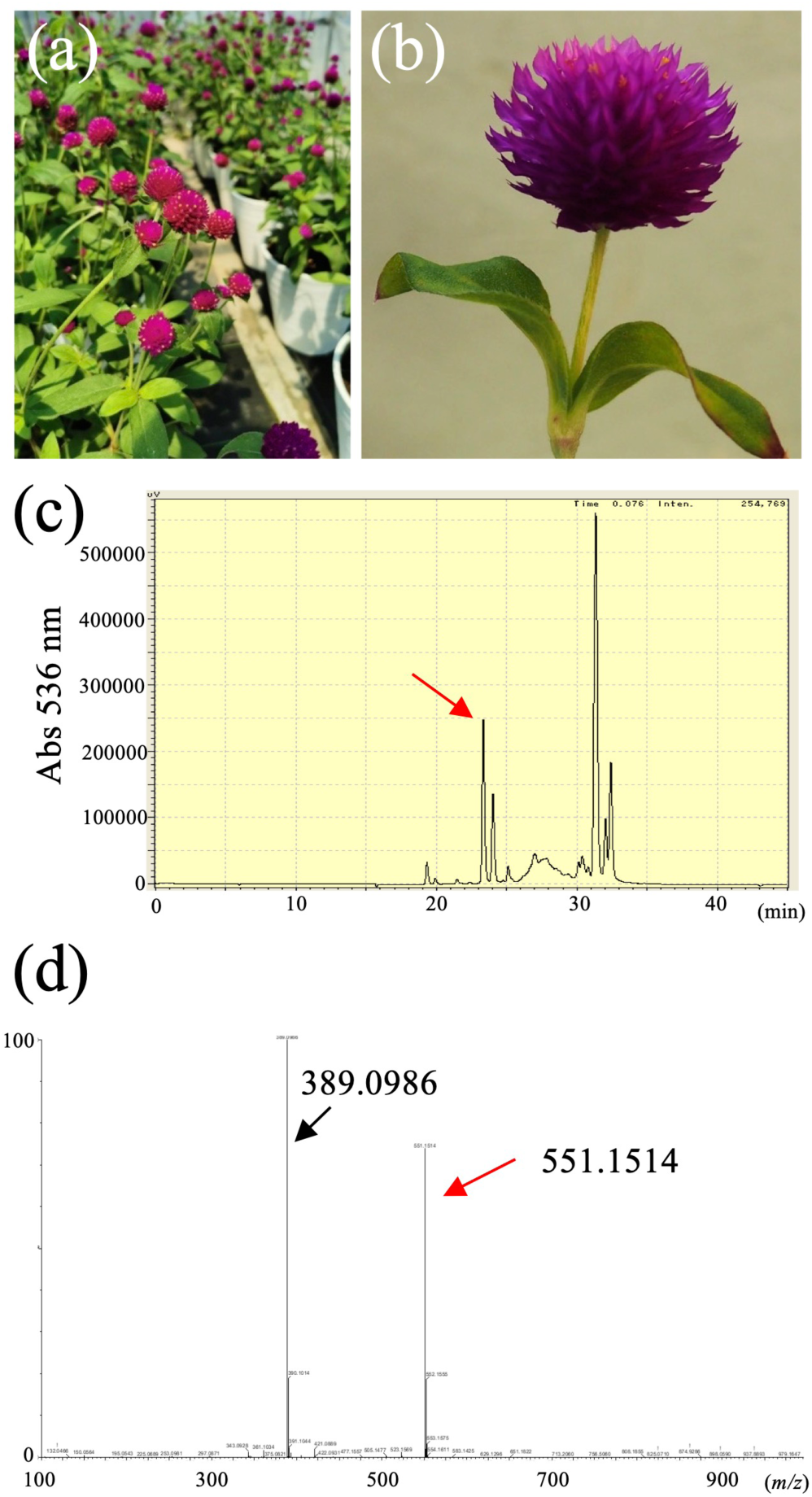
Pigment analysis of *Gomphrena globosa*. (a, **b)** Photographs of *G. globosa* plants **(a)** and flower bracts **(b)**. **(c)** HPLC chromatogram of extracts from *G. globosa* flower bracts. The red arrow indicates gomphrenin I. The horizontal axis represents retention time (min), and the vertical axis represents signal intensity (µV). **(d)** Mass spectrum of the HPLC-eluted pigment corresponding to the arrow in **(c)**. The spectrum was obtained from *G. globosa* flower bracts. The red peak corresponds to gomphrenin I. The horizontal axis indicates mass-to-charge ratio (*m/z*), and the vertical axis indicates relative abundance. Arrows denote betanidin fragments derived from gomphrenin I.

The enzymatic activity of GgcDOPA6GT was assessed using a transient expression assay in *N. benthamiana* leaves. To do so, an expression plasmid carrying *GgcDOPA6GT* (Figure S4) was introduced into *A. tumefaciens*, and the resulting strains were co-infiltrated with *CqCYP76AD1-1* and *CqDODA-1* into *N. benthamiana* leaves. Red pigmentation developed in the infiltrated tissues (Figure 5a). RT-PCR analysis confirmed expression of all transgenes in these red-colored leaves (Figure 5b). Pigments extracted from the infiltrated tissues were subsequently analyzed by HPLC, which showed that GgcDOPA6GT predominantly produced gomphrenin I (Figure 5c).

**Figure 5.**
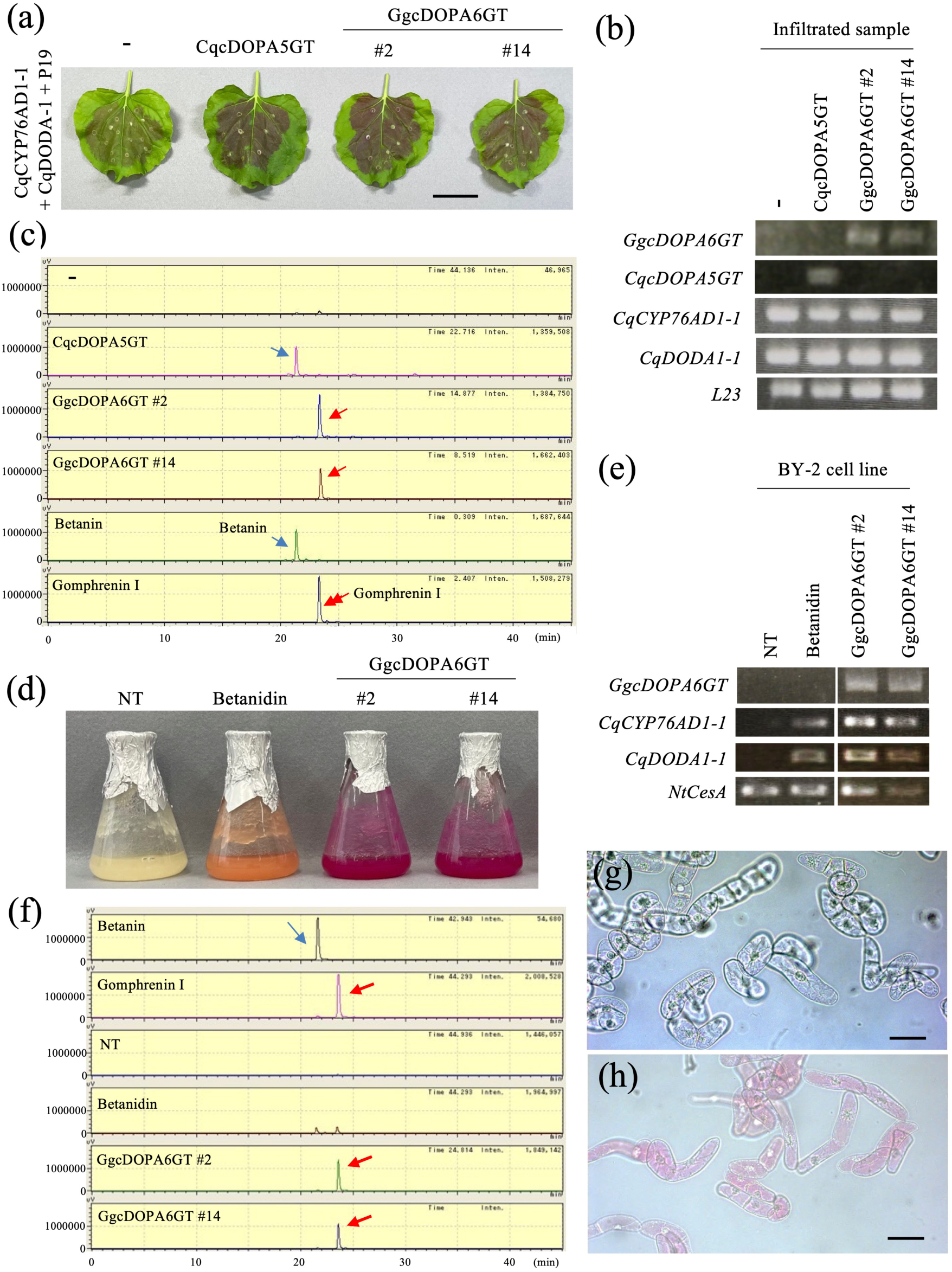
Functional characterization of *GgcDOPA6GT* from *Gomphrena globosa*. **(a)** Recombinant expression of *GgcDOPA6GT* and *CqcDOPA5GT* in *N. benthamiana* leaves. *Agrobacterium* strains carrying *GgcDOPA6GT* or *CqcDOPA5GT* were co-infiltrated with CqCYP76AD1-1, CqDODA-1, and P19. The minus sign (−) indicates samples co-infiltrated with CqCYP76AD1-1, CqDODA-1, and P19 only. #2 and #14 indicate independent *GgcDOPA6GT* clones. Bar, 4 cm. **(b)** RT-PCR analysis of transgene expression in infiltrated *N. benthamiana* leaves. *L23* was used as an internal control. **(c)** HPLC chromatogram of pigments extracted from infiltrated *N. benthamiana* leaves. Betanin and gomphrenin I standards are shown for comparison. Blue and red arrows indicate the peaks corresponding to betanin and gomphrenin I, respectively. The horizontal axis shows retention time (min), and the vertical axis shows signal intensity (µV). **(d)** Photographs of transformed tobacco BY-2 cell lines two weeks following subculturing. GgcDOPA6GT indicates lines expressing GgcDOPA6GT together with CqCYP76AD1-1, and CqDODA-1. “Betanidin” denotes lines expressing only CqCYP76AD1-1 and CqDODA-1. “NT” indicates the non-transgenic BY-2 cell line. #2 and #14 represent independent transformants. **(e)** RT-PCR analysis of transgene expression, with NtCesA as the internal control. **(f)** HPLC chromatograms of pigments from transformed lines. Blue and red arrows marking betanin and gomphrenin I, respectively. The horizontal axis shows retention time (min), and the vertical axis shows signal intensity (µV). **(g, h)** Microscopy images of NT cells **(g)** and gomphrenin I–producing line (*GgcDOPA6GT* #2) **(h)**. Bars, 100 µm.

To further characterize its enzymatic activity, we next established a BY-2 cell line that also expressed *GgcDOPA6GT*. The transformed *Agrobacterium* strain harboring the *GgcDOPA6GT* construct was introduced into tobacco BY-2 cells. A stable *GgcDOPA6GT*-expressing BY-2 cell line containing *CqCYP76AD1-1*, *CqDODA-1*, and *GgcDOPA6GT* was thus obtained (Figures 5d and S8). After confirming transgene expression by RT-PCR (Figure 5e), a highly pigmented line was selected and maintained in liquid culture. The *GgcDOPA6GT*-expressing lines showed vivid red coloration (Figure 5d). Subsequent HPLC analysis indicated that these lines produced gomphrenin I as the major pigment (Figure 5f), with concentrations of 46.66 ± 3.15 µM. Similar pigment accumulation profiles were observed across additional independent lines (Figure S8). In these lines, gomphrenin I accumulated in vacuoles, similar to what was observed in the BacDOPA5/6GT-expressing lines (Figure 5g and h). These findings suggest that GgcDOPA6GT, unlike BacDOPA5/6GT from *B. alba*, exhibits specific cDOPA6GT activity and catalyzes gomphrenin I biosynthesis via cDOPA 6-O-glucoside.

### Phylogenetic analysis of the cDOPA glucosyltransferase family

In this study, we successfully isolated genes encoding enzymes with cDOPA6GT activity from *B. alba* and *G. globosa* (i.e., BacDOPA5/6GTs and GgcDOPA6GT). To clarify how these enzymes relate phylogenetically to previously characterized cDOPA5GTs, we conducted phylogenetic analysis using their amino acid sequences (Table S4). The resulting tree showed that GgcDOPA6GT grouped within a clade composed of the Amaranthaceae members, whereas BacDOPA5/6GTs were placed in a distinct clade (Figure 6). Since only a limited number of enzymes in each clade have been functionally characterized, the functional assignments of these clades should be interpreted with caution. In addition, sequences from non-Caryophyllales species were included to provide broader phylogenetic context and do not necessarily represent cDOPA-related enzymatic functions.

**Figure 6.**
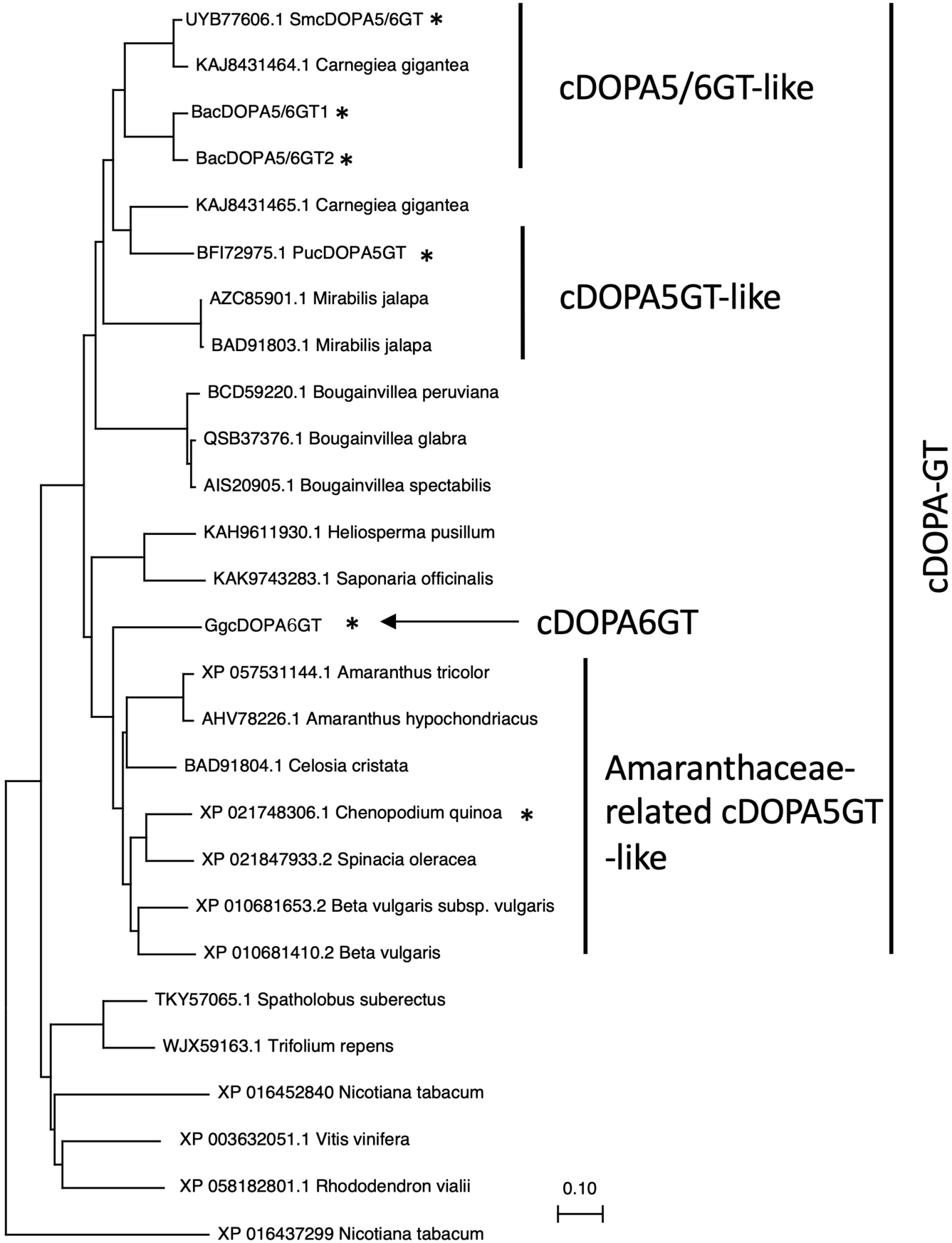
Molecular phylogenetic tree of selected cyclo-DOPA O-glucosyltransferases (cDOPA-GTs). Amino acid sequences were aligned using MUSCLE, and a phylogenetic tree was constructed with the maximum-likelihood method in MEGA11. Bootstrap support from 5,000 replicates is shown at branch nodes. The scale bar represents 0.1 substitutions per site. Asterisks mark the genes whose cDOPA-GT activities were experimentally evaluated. “cDOPA-GT” indicates the main clade, and “cDOPA5GT-like,” “cDOPA5/6GT-like,” and “Amaranthaceae-related cDOPA5GT-like” denote subclusters. Homologous sequences from additional plant species are listed in Table S4. Functional classifications are based on currently characterized enzymes and should be interpreted cautiously. Sequences from non-Caryophyllales species were included to provide phylogenetic context.

To evaluate whether additional plant species possess cDOPA6GT activity, we examined the subclade containing other BacDOPA5/6GT protein. This subclade also included PucDOPA5GT from *Portulaca umbraticola* (Acc. No.: BFI72975.1) and SmcDOPA5/6GT from *Selenicereus monacanthus* (Acc. No.: UYB77606.1). The amino acid sequence identities of PucDOPA5GT and SmcDOPA5/6GT with BacDOPA5/6GT1 were 69% and 75%, respectively (Table S1).

Next, to assess the enzymatic activities of PucDOPA5GT and SmcDOPA5/6GT, expression plasmids were generated for each gene (Figure S4) and introduced into *A. tumefaciens*. The transformed *Agrobacterium* strains harboring these constructs, together with CqCYP76AD1-1 and CqDODA-1, were co-infiltrated into *N. benthamiana* leaves. Red pigmentation was developed in leaves expressing either PucDOPA5GT or SmcDOPA5/6GT (Figure S11). HPLC analysis of pigments extracted from the infiltrated tissues showed that PucDOPA5GT produced only betanin, whereas SmcDOPA5/6GT yielded both betanin and gomphrenin I (Figure S11c). These findings highlight the divergent evolution of cDOPA-GT catalytic functions in betalain-producing species.

### Structural and functional analyses of cDOPA glucosyltransferase active sites

This study revealed that the cDOPA glucosyltransferase (cDOPA-GT) family comprises three putative functional types: cDOPA5GT-like, dual cDOPA5/6GT-like, and cDOPA6GT-like enzymes. To identify residues required for catalysis, we selected residues for mutagenesis based on their sequence conservation among cDOPA-GTs, AlphaFold-based structural modeling (Figure S12), and their predicted roles in catalysis or substrate binding within the active site. Two conserved histidines in CqcDOPA5GT, BacDOPA5/6GT1, and GgcDOPA6GT were replaced with alanine (Figures 7a and S7). Site-directed mutagenesis was performed to generate the indicated variants, and their enzymatic activities were experimentally evaluated using the transient expression system in *N. benthamiana.* When these substitutions were introduced into *BacDOPA5/6GT1* and *GgcDOPA6GT*, red pigmentation in *N. benthamiana* was strongly reduced, demonstrating that both histidines are critical for activity (Figures 7b, c). In contrast, the two substitutions produced different outcomes in CqcDOPA5GT: the N-terminal variant (H23A) lowered pigmentation, whereas the C-terminal variant (H369A) caused a pronounced increase in pigment accumulation.

**Figure 7.**
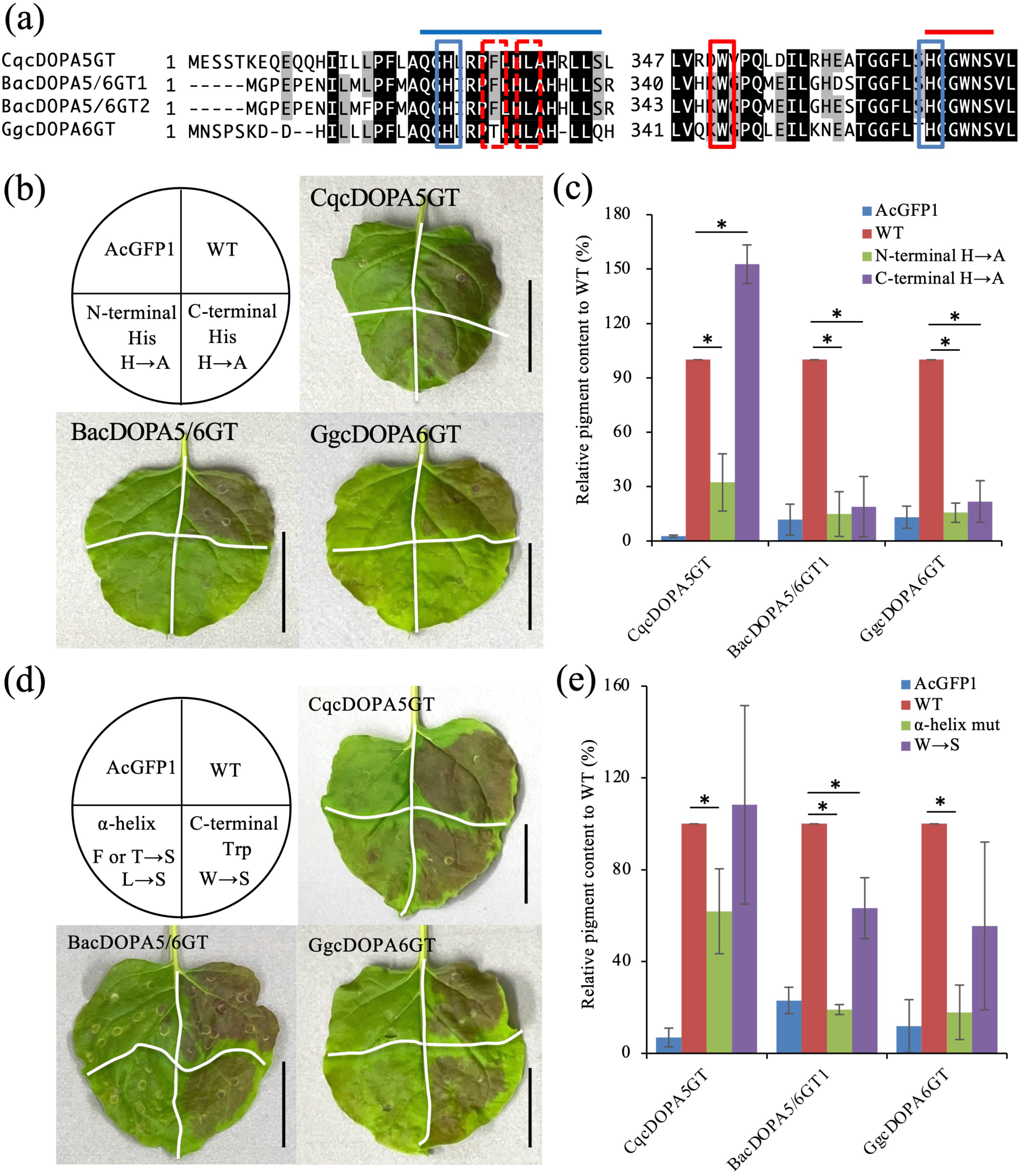
Evaluation of putative active-site residues in the cDOPA-glucosyltransferase family. **(a)** Alignment of deduced amino acid sequences of CqcDOPA5GT, BacDOPA5/6GTs, and GgcDOPA6GT. Blue boxes indicate residues predicted to participate in catalysis. A dashed red box and a solid red box highlight residues suggested by 3D simulation to influence activity despite lying outside the catalytic site. The α-helical region and the conserved UGT motif are indicated by blue and red lines, respectively. **(b)** Recombinant expression of alanine-substituted His mutants in *N. benthamiana* leaves. *Agrobacterium* carrying wild-type and mutants cDOPA-GT constructs were co-infiltrated with CqCYP76AD1-1, CqDODA-1, and P19. The N-terminal and C-terminal His residues correspond to the black and red asterisks in (a), respectively. AcGFP1 served as the negative control. Bar, 4 cm. **(c)** Relative pigment production of His→Ala mutants. The vertical axis shows pigment accumulation relative to WT (%). Error bars represent mean ± SD (n = 3). \**p* < 0.05 vs. WT (Student’s *t*-test). **(d)** Recombinant expression of Ser-substituted mutants targeting the α-helix or the conserved C-terminal Trp residue, corresponding to the sites boxed in (a). **(e)** Relative pigment production of Ser-substituted mutants, The vertical axis shows the pigment accumulation relative to WT (%). Error bars represent mean ± SD (n = 3). \**p* < 0.05 vs. WT (Student’s *t*-test).

Structural modeling of BacDOPA5/6GT1, guided by the crystal structure of TcCGT1, a C-glycosyltransferase from *Trollius chinensis* (PDB: 6JTD), identified an α-helical region together with a nearby tryptophan positioned next to the catalytic pocket (Figure S13). Residues within this α-helical region were selected based on their proximity to the active site and mutated to serine to disrupt local side-chain interactions while preserving overall structural integrity. When this α-helix was disrupted, activity was abolished in BacDOPA5/6GT1 (F22S/L25S) and in GgcDOPA6GT (T24S/L27S), whereas the corresponding CqcDOPA5GT mutant retained roughly 60% activity (Figures 7d, e).

Substitution of the conserved tryptophan similarly reduced pigment production in BacDOPA5/6GT1 (W344S) and GgcDOPA6GT (W345S), although the equivalent CqcDOPA5GT variant (W351S) showed only a minor reduction (Figures 7d, e). Taken together, these observations indicate that the α-helical region adjacent to the catalytic site is essential for BacDOPA5/6GT1 and GgcDOPA6GT, while it contributes only partially to CqcDOPA5GT function.

### Comparative physicochemical property of gomphrenin I and betanin

We obtained gomphrenin I in sufficient quantity and purity for nuclear magnetic resonance (NMR) structural determination. Gomphrenin I is a positional isomer of betanin with the same molecular weight; however, it shows a distinct absorption maximum at 543 nm, compared with 536 nm for betanin (Belhadj Slimen et al., 2017), and it also displays a different retention time on reversed-phase HPLC (Figure 1e). These features suggest that gomphrenin I may exhibit physical properties that diverge from those of betanin. Therefore, we compared the thermal stability of gomphrenin I and betanin.

To evaluate the thermal stability of betanin and gomphrenin I, heat treatments were applied for 3 h using a thermal cycler. Under these conditions, gomphrenin I remained significantly more stable than betanin at temperatures above 40°C (Figure 8a–c, Figure S14). Thermal treatment was then examined by NMR spectroscopy. Samples were incubated for 15 min at the designated temperatures, and the one-dimensional NMR spectra were recorded after every step. The temperature was increased sequentially from 26°C to 65°C, initially by 4°C and then in 5°C increments. Betanin maintained its native structure up to 30°C, but structural alterations were evident above 35°C (Figure 8d). In contrast, gomphrenin I retained its structure up to 40°C, with detectable changes emerging at 45°C and higher (Figure 8e). These results demonstrate that gomphrenin I displays greater thermal stability than betanin, and neither pigment recovered its original structure after heat-induced denaturation. To explore a possible structural basis for this difference, computational chemistry analyses were conducted. Molecular dynamics (MD) simulations and density functional theory (DFT) calculations indicated that gomphrenin I can form a bridging intramolecular hydrogen bond between the glucose moiety and the betalamic acid moiety, a feature not predicted for betanin (Figure S15). The presence of this intramolecular hydrogen bond may therefore contribute to the enhanced structural stability of gomphrenin I relative to betanin.

**Figure 8.**
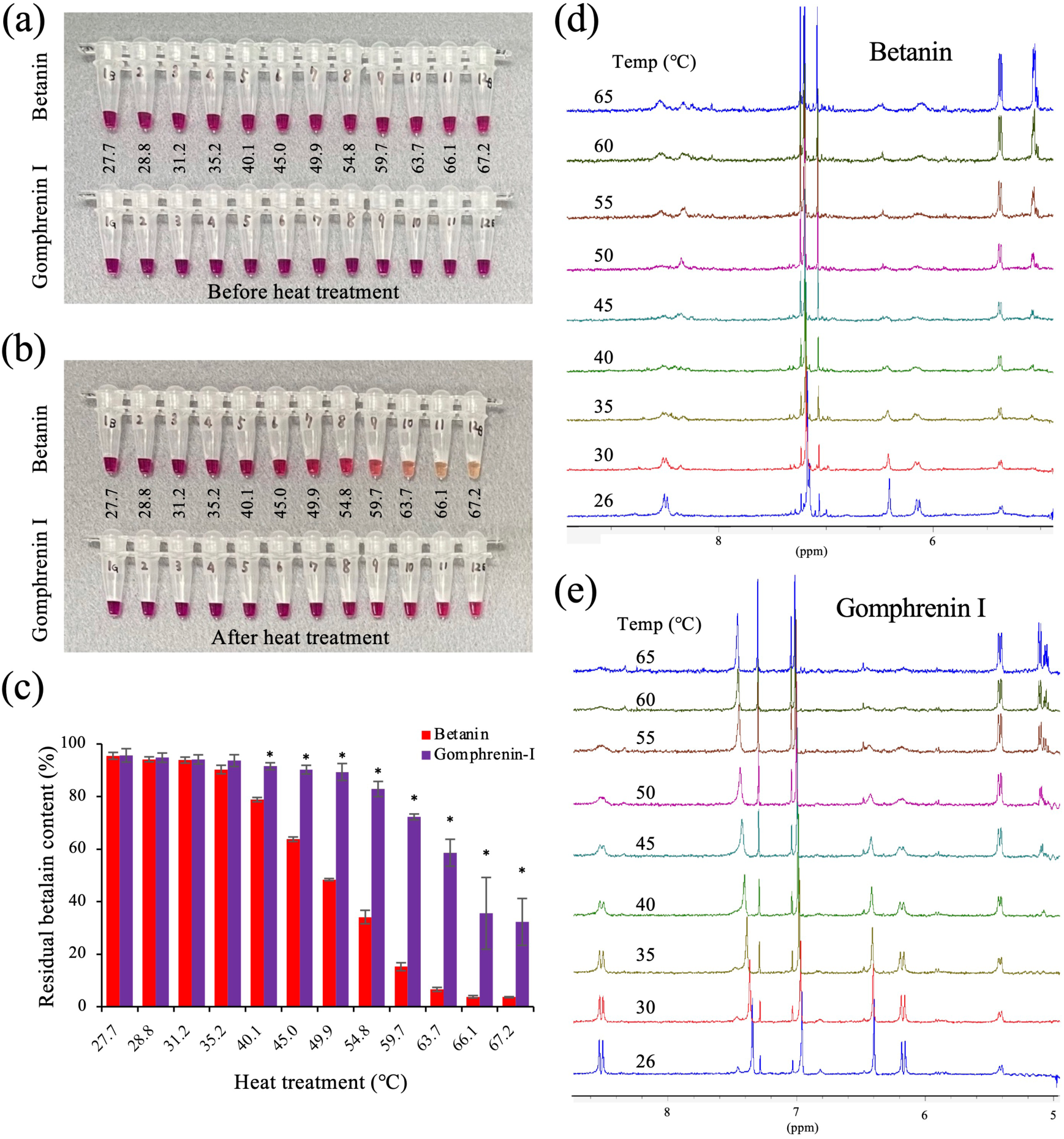
Thermal stability of gomphrenin I and betanin. (a, **b)** Photographs of gomphrenin I and betanin solutions before **(a)** and after **(b)** heat treatment at the temperatures indicated (℃). **(c)** Thermal stability of both pigments. The vertical axis shows residual pigment content (%) after heating, and the horizontal axis shows treatment temperature (℃). Error bars indicate mean ± SD (n = 3); \**p* < 0.05 vs. betanin. **(d, e)** ^1^H NMR spectra of heated betanin **(d)** and gomphrenin I **(e)**, with temperatures indicated.

## Discussion

In this study, we identified and characterized cDOPA6GT enzymes involved in gomphrenin I biosynthesis in B. alba and G. globosa, revealing a previously unrecognized biosynthetic route. We cannot exclude the possibility that additional UGTs contribute to betalain glycosylation in planta; however, the candidates analyzed in this study are likely to represent the relevant enzymes based on sequence similarity and expression profiles.

Consistent with previous studies, the biosynthesis of gomphrenin I, which carries a glucose moiety at the C6 position of betanidin, is catalyzed by betanidin 6-O-glucosyltransferase (B6GT) (Vogt et al., 2002). In contrast, two alternative routes are known for betanin biosynthesis: one involves glucosylation of betanidin by betanidin 5-O-glucosyltransferase (B5GT), and the other involves glucosylation of cyclo-DOPA by cyclo-DOPA 5-O-glucosyltransferase (cDOPA5GT) (Figure S1). Our results show that BacDOPA5/6GTs and GgcDOPA6GT, isolated as orthologs of CqcDOPA5GT, possess cDOPA6GT activity involved in gomphrenin I biosynthesis. These findings indicate that, similar to betanin biosynthesis, gomphrenin I biosynthesis proceeds through two distinct pathways: one mediated by B6GT and the other by cDOPA6GT (Figure 1a). Although both pathways appear to be present, it remains unclear which pathway is predominant in each species or tissue. The relative contribution of the B6GT-mediated and cDOPA6GT-mediated routes may vary depending on species, developmental stage, or tissue type. Further quantitative analyses will be required to determine the relative importance of these pathways in plants.

We further defined *three* functional classes of cyclo-DOPA O-glucosyltransferases: those with cDOPA5GT activity (e.g., CqcDOPA5GT), those with dual cDOPA5/6GT activity (e.g., BacDOPA5/6GT), and those with dedicated cDOPA6GT activity (e.g., GgcDOPA6GT). These enzymes share roughly 60% amino acid identity and cluster within the cDOPA-GT clade. Within this clade, cDOPA5GT-like proteins span the major lineage, whereas the cDOPA5/6GTs-type protein form a distinct subclade.

GgcDOPA6GT did not group within this subclade but instead occupied a neighboring cluster containing Amaranthaceae-type cDOPA5GTs (Figure 6). This pattern suggests that cDOPA6GT activity arose independently in GgcDOPA6GT and the cDOPA5/6GT-type proteins through distinct evolutionary trajectories. Additional characterization of cDOPA-GTs from other gomphrenin I-producing species will be necessary to resolve the evolutionary origins of these catalytic functions.

Our mutational analysis showed that the N-terminal His residue is essential for cDOPA-GT catalytic activity, likely forming part of the active site. The C-terminal His was also required in BacDOPA5/6GT and GgcDOPA6GT, whereas its substitution in CqcDOPA5GT unexpectedly increased product accumulation, indicating that this residue is less critical in cDOPA5GT (Figure 7). We also identified an α-helix adjacent to the catalytic pocket that strongly influence enzyme activity, particularly in cDOPA5/6GT1 and GgcDOPA6GT. Disrupting the hydrophobic interface formed by this α-helix and the adjacent β-sheet markedly reduced activity (Figure S13). Such distal structural features can modulate enzyme function via long-range or allosteric mechanisms consistent with the “butterfly effect” (Ohmae et al., 1996). The reduced activity observed in the α-helix mutants likely reflects perturbation of the substrate-binding pocket caused by the collapse of this structural element. However, it should be noted that the effects of mutations on protein folding were not directly assessed in this study. Therefore, we cannot exclude the possibility that some of the observed changes in enzymatic activity may be partially influenced by alterations in protein stability or folding. Attempts to pinpoint residues determining product specificity were inconclusive: structural predictions using AlphaFold and related tools revealed no clear active-site differences among cDOPA-GTs, and none of the mutants showed detectable changes in specificity. These results indicate that the observed differences reflect product specificity among cDOPA-GTs rather than substrate specificity, as all enzymes utilize cyclo-DOPA as a common substrate. Co-crystal structures of cDOPA-GTs with their substrates will be required to resolve these determinants and to clarify the long- range structural coupling suggested here.

Although BacDOPA5/6GTs catalyzed the production of both betanin and gomphrenin I in heterologous systems (Figure 2 and 3), only gomphrenin-type pigments are detected in *B. alba*, and betanin has not been detected (Figure 1e). This discrepancy suggests that biosynthetic processes occurring in planta differ from those observed in heterologous plant expression systems. Several explanations are plausible *B. alba* may encode an additional, as yet unidentified enzyme with exclusive cDOPA6GT activity, analogous to GgcDOPA6GT. Alternatively, the B6GT identified here, or another uncharacterized B6GT, may act preferentially at C6 *in vivo*. It is also possible that endogenous cofactor or protein partners in *B. alba* shift the activity of BacDOPA5/6GT toward cDOPA6GT. At present, these mechanisms cannot be tested directly because genomic resources for *B. alba* are unavailable and genetic manipulation of betalain-producing plants, including stable transformation, remains technically challenging. Future genomic and functional analyses will be essential to explain the divergence between heterologous and native enzymatic activities.

In this study, the comparison between gomphrenin I and betanin was conducted to examine the impact of glycosylation position on pigment stability, particularly in the context of the newly identified cDOPA 6-O-glucosylation pathway. Thermal stability analysis revealed that gomphrenin I retained its structural integrity at higher temperatures than betanin (Figures 8 and S14). Betalains have been reported to exhibit strong antioxidant activity and are considered to play roles in protecting plants against abiotic stresses including high light, drought, and temperature stress (Grotewold, 2006; Gould, 2004). In addition, recent reports have described various physiological activities of betalain pigments (Allegra et al., 2014; Nowacki et al., 2015; Imamura et al., 2022). Thermal stability under physiological conditions (approximately 37°C) is likely important for these activities to be exerted effectively. In NMR-based thermal stability assays, betanin exhibited structural alteration at 30°C, whereas gomphrenin I maintained its structure up to 40°C. It should be noted that the thermal stability assessed by UV–Vis spectroscopy and NMR reflects different aspects of pigment stability. While spectrophotometric measurements primarily monitor changes in chromophore integrity and pigment absorbance, NMR analysis provides information on the preservation of molecular structure. Therefore, differences between these approaches may arise from variations in sensitivity to partial degradation or structural rearrangements. Greater thermal stability of gomphrenin I suggests that it may retain its physiological functions more effectively than betanin under biological conditions. Beyond potential effects on physiological activity, differences in thermal stability may also influence pigment utilization in plants. It is therefore plausible that the greater thermal stability of gomphrenin I contributes to its deployment in certain Caryophyllales species, including tropical plants such as *B. alba* and long-lasting ornamental flowers like *G. globosa.* It is also notable that betanidin itself was not detected in control systems, suggesting that this aglycone is rapidly converted into glycosylated derivatives *in vivo* rather than accumulating as a stable intermediate. This observation supports the idea that endogenous glycosyltransferase activities contribute to the low-level accumulation of betalain pigments in control samples. However, ecological or physiological studies are required to test this possibility.

Further, computational analyses provided a plausible structural basis for the enhanced thermal stability of gomphrenin I. Betacyanins are generally thermally unstable, often degrading via hydrolysis of the iminium bond (Silva et al., 2022). However, under our experimental conditions, betalamic acid, which would be expected as a product of this degradation pathway, was not detected (Figure S16). This suggests that the thermal degradation observed in this study proceeds via a mechanism distinct from previously reported pathways. Our calculations suggested that gomphrenin I may form a unique bridging intramolecular hydrogen bond between the 6-O- glucosyl moiety and the betalamic acid moiety (Figure S15a). This hydrogen bond, which was not predicted for betanin (Figure S15b), links two portions of the molecule that would otherwise be cleaved during thermal degradation. It is therefore likely that this local stabilization, preventing cleavage, contributes to the observed increase in overall thermal stability. Thus, the position of glucosylation, and its capacity to support this intramolecular bridge, appears to be a key structural determinant for the distinct thermal stability observed between these two betacyanin isomers. However, this interaction may be influenced by solvent effects and conformational dynamics in solution and therefore may not always be favored under physiological conditions.

In this study, we identified BacDOPA5/6GTs from *B. alba* and GgcDOPA6GT from *G. globosa* as enzymes involved in gomphrenin I biosynthesis. Similar examples of convergent biosynthetic pathways have been reported in plant specialized metabolism (Grotewold, 2006), where distinct enzymatic routes can produce identical end products, such as in flavonoid and alkaloid biosynthesis. Our results indicate that gomphrenin I formation can proceed not only via the previously described B6GT-mediated route but also through an alternative cDOPA6GT-mediated pathway. This pathway-level diversification provides insight into the molecular basis underlying the structural diversity of betacyanins and clarifies how distinct glucosylation steps contribute to branching within the betacyanin biosynthetic network. In addition, the establishment of a BY-2 cell-based production system enabled direct comparison of the physicochemical properties of gomphrenin I and its positional isomer, betanin, offering a useful experimental framework for future studies of structure–function relationships in betalain pigments. Together, these findings advance our understanding of the enzymatic and biochemical mechanisms shaping betacyanin diversity and provide a foundation for exploring their functional significance in plant metabolism.

## Acknowledgments

The authors thank Akiko Mizuno and Megumi Sakuda for technical assistance. The authors also thank Enago (www.enago.jp) for English language review. The authors also acknowledge experimental support from the Advanced Research Infrastructure for Materials and Nanotechnology in Japan (ARIM; MEXT, Japan). This work was partially supported by KAKENHI Grants (Nos. 20K06049, 21H05032, and 25H00431 [to S.O.], and 23K05228, and 26K08789 [to T.I.]).

## Author contributions

MM conceived this study. TI and MM designed the experiments. TI, RS, and MM isolated the genes used in this study. KM, TS, MY, and HT performed RNA-seq analyses. TI, RS, MM, and NM conducted experiments to evaluate the candidate gene activity using *N. benthamiana*. TI, RS, MM and NM conducted HPLC analyses. MM established the betacyanin production system using BY-2 cells. RS and MM purified gomphrenin I and betanin. AM performed LC-MS analysis. RS carried out thermal stability assay. SO carried out NMR analysis and simulation analyses. TY performed MD and DFT calculations. TI, SO, TY, and MM wrote the manuscript. All authors have read and approved the final version of the manuscript.

## Data Availability Statement

The data that support the findings of this study are included in this published article and its Supporting Information files. RNA-seq data have been deposited in the DDBJ Sequence Read Archive under accession numbers DRX780510 and DRX780511 (*B. alb*a) and DRX780505–DRX780509 (*G. globosa*).

## Supporting Information

Additional Supporting Information may be found online in the Supporting Information section.

