## Supplemental Figures 1-20 for "A cyclo-DOPA 6-O-glucosyltransferase-mediated route for gomphrenin I biosynthesis in *Basella alba* and *Gomphrena globosa*"

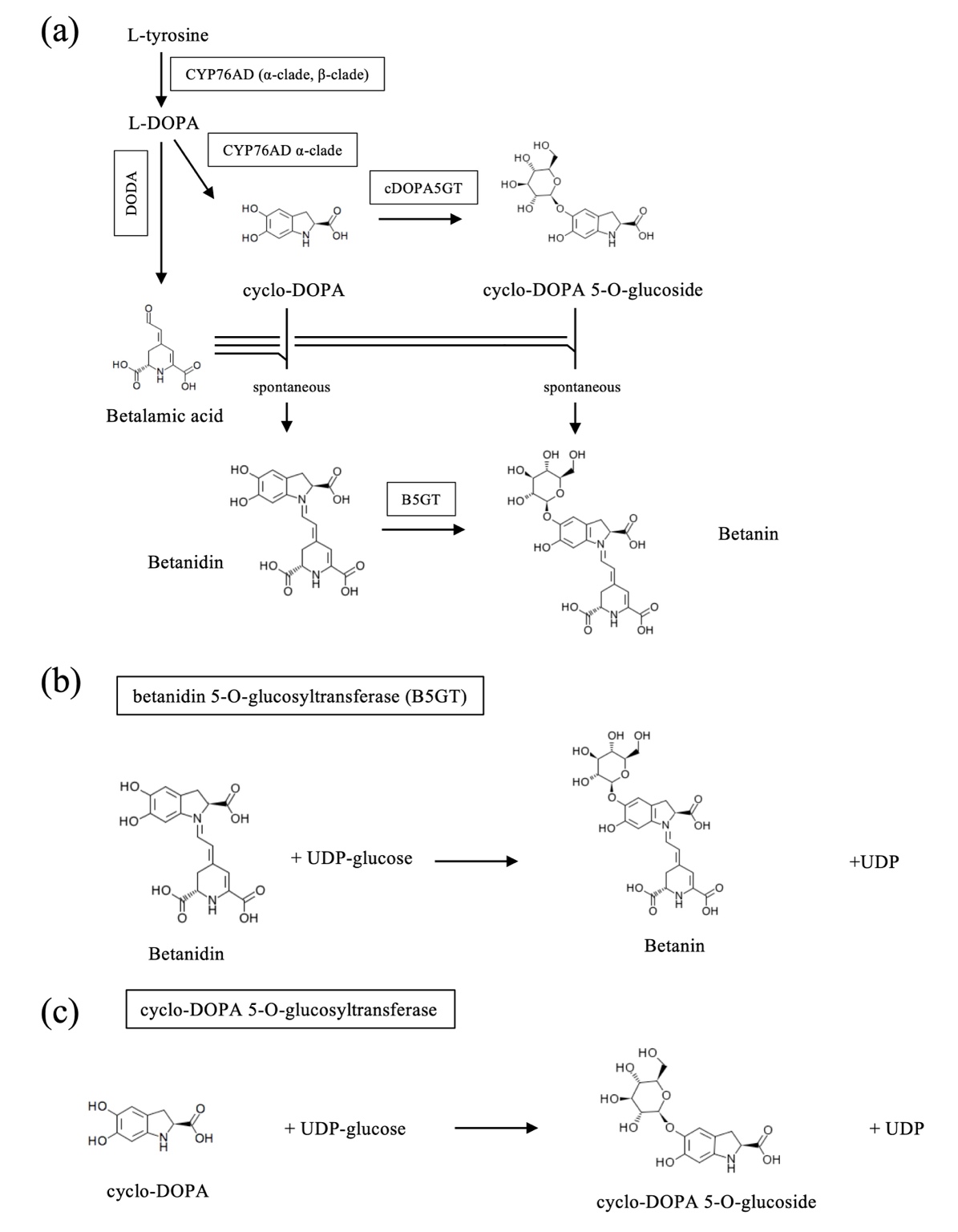

**Figure S1. Schematic representation of the betanin biosynthetic pathway and the cDOPA5GT-catalyzed reaction. (a)** Schematic diagram of the betanin biosynthetic pathway. Boxes indicate betalain biosynthetic enzymes. CYP76AD1, cytochrome P450 76AD1; DODA1, DOPA 4,5-dioxygenase 1; cDOPA5GT, cyclo-DOPA 5-O-glucosyltransferase; B5GT, betanidin 5-O-glucosyltransferase. **(b, c)** Schematic representation of the reaction for betanin **(b)**, and cyclo-DOPA-5-O-glucuronylglucoside **(c)**.

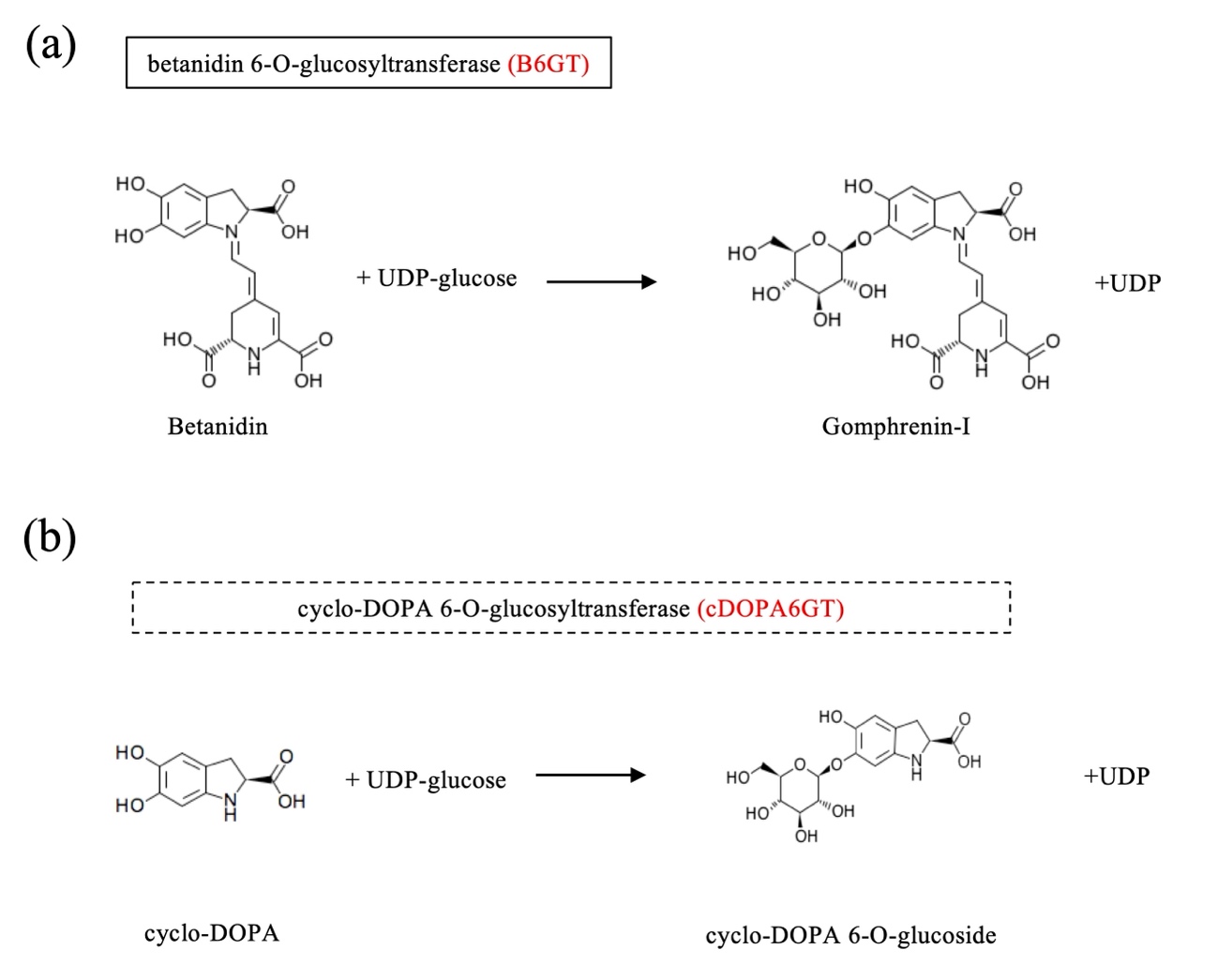

**Figure S2. Schematic representation of reaction for Gomphrenin I(a), and cyclo-DOPA-6-O-glucuronylglucoside (b).** Boxes and dashed boxes indicate identified and predicted enzymes.

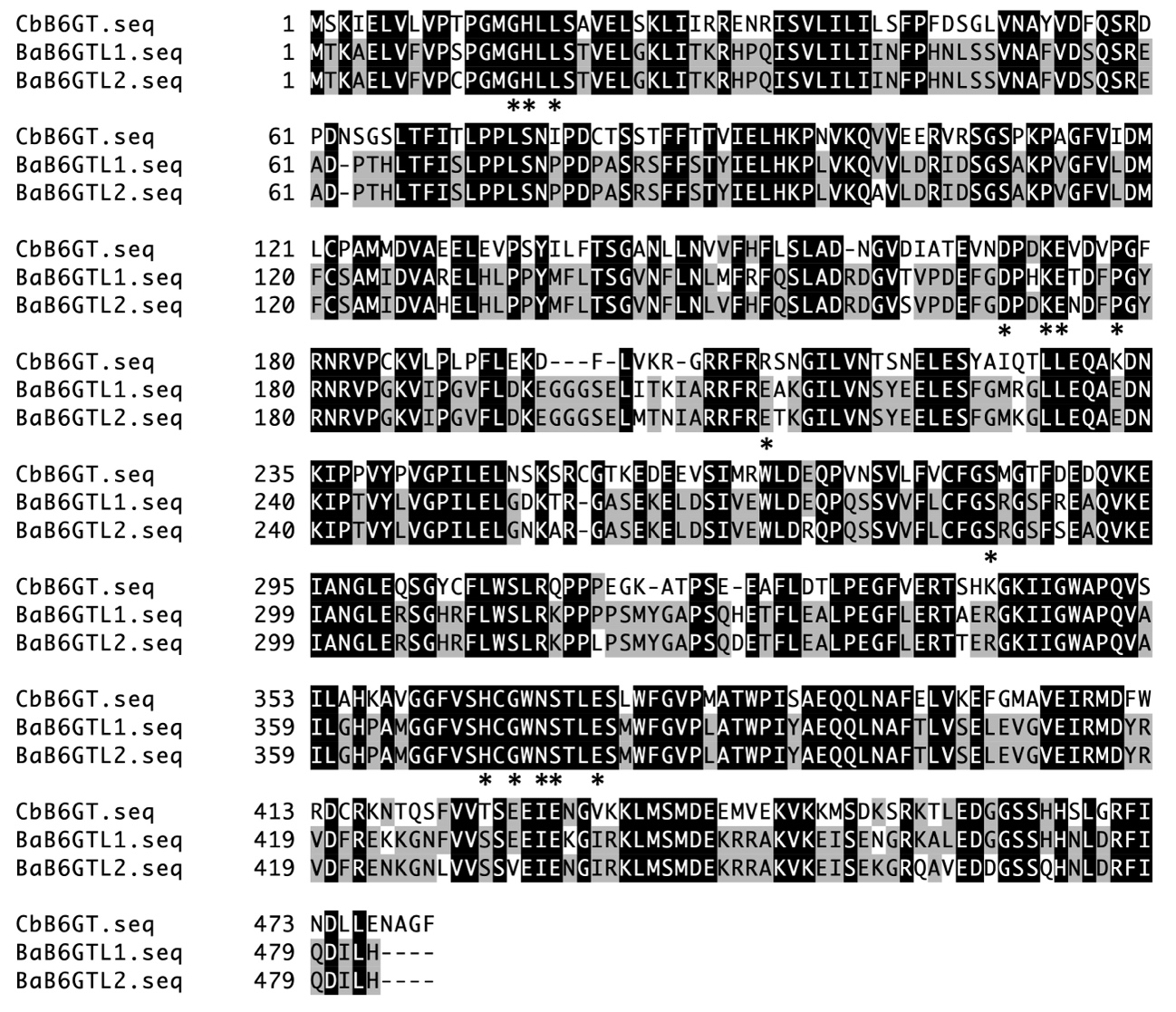

**Figure S3. Alignment of the deduced amino acid sequences of CbB6GT, BaB6GT1, and BaB6GT2.** Asterisks indicate amino acid residues expected to form the active site of CbB6GT.

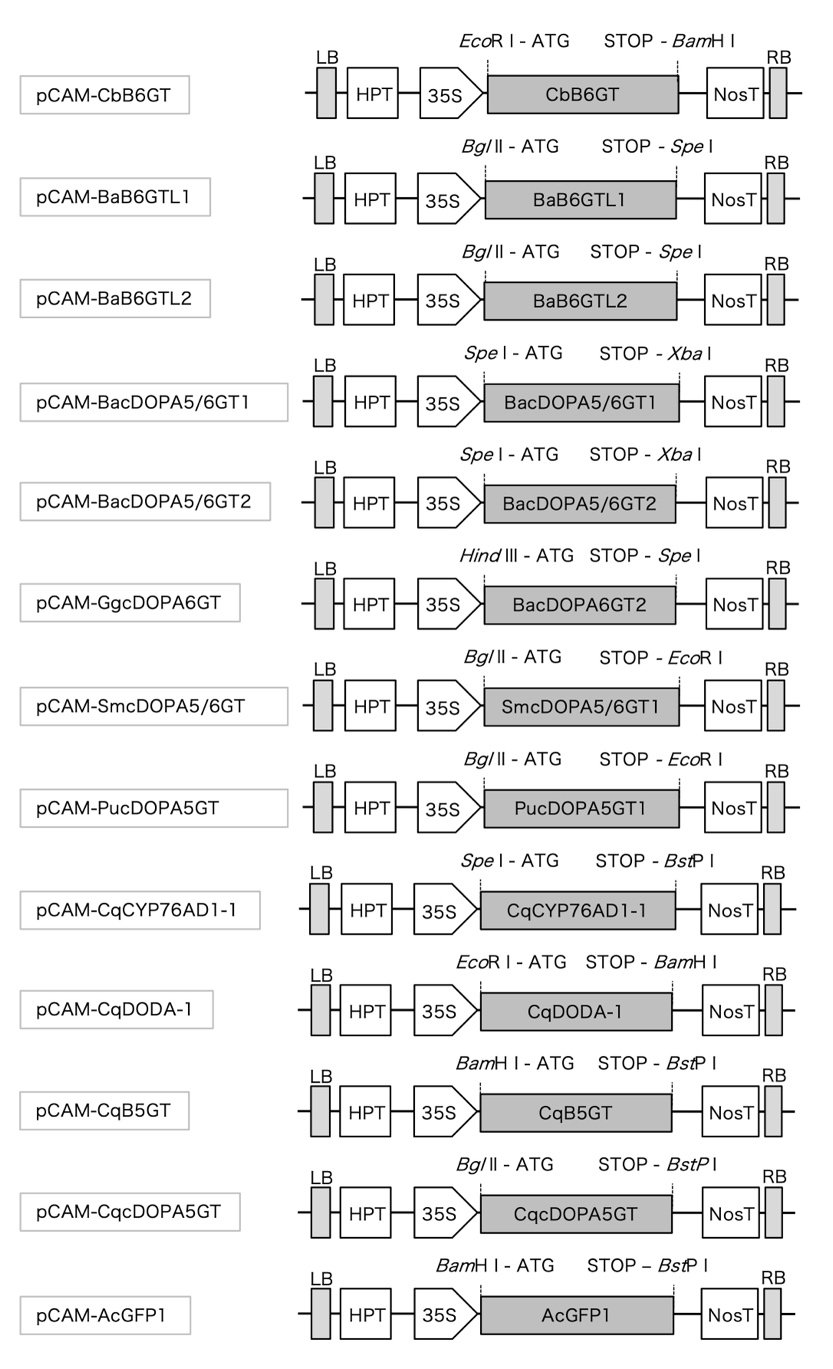

**Figure S4. Schematic representations of the plant expression vector.** CbB6GT, *CbB6GT* CDS; BaB6GTL1, *BaB6GTL1* CDS; BaB6GTL2, *BaB6GTL2* CDS; BacDOPA5/6GT1, *BacDOPA5/6GT1* CDS; BacDOPA5/6GT2, *BacDOPA5/6GT2* CDS; GgcDOPA6GT, *GgcDOPA6GT* CDS; SmcDOPA5/6GT, *SmcDOPA5/6GT* CDS; PucDOPA5GT, *PucDOPA5GT* CDS; CqcDOPA5GT, *CqcDOPA5GT* CDS; CqB5GT, *CqB5GT* CDS; CqCYP76AD1-1, *CqCYP76AD1-1* CDS; CqDODA-1, *CqDODA-1* CDS; AcGFP1, *AcGFP1* CDS; 35S, CaMV 35S promoter; NosT, *nopaline synthase* terminator; 35S-T, 35S terminator; RB, right border; LB, left border; HPT, *hygromycin phosphotransferase* expression cassette; ATG, start codon; STOP, stop codon. Abbreviations for species: Ba, *Basella alba*; Cq, *Chenopodium quinoa*; Cb, *Cleretum bellidiforme*; Gg, *Gomphrena globosa*; Sm, *Selenicereus monacanthus*; Pu, *Portulaca umbraticola.*

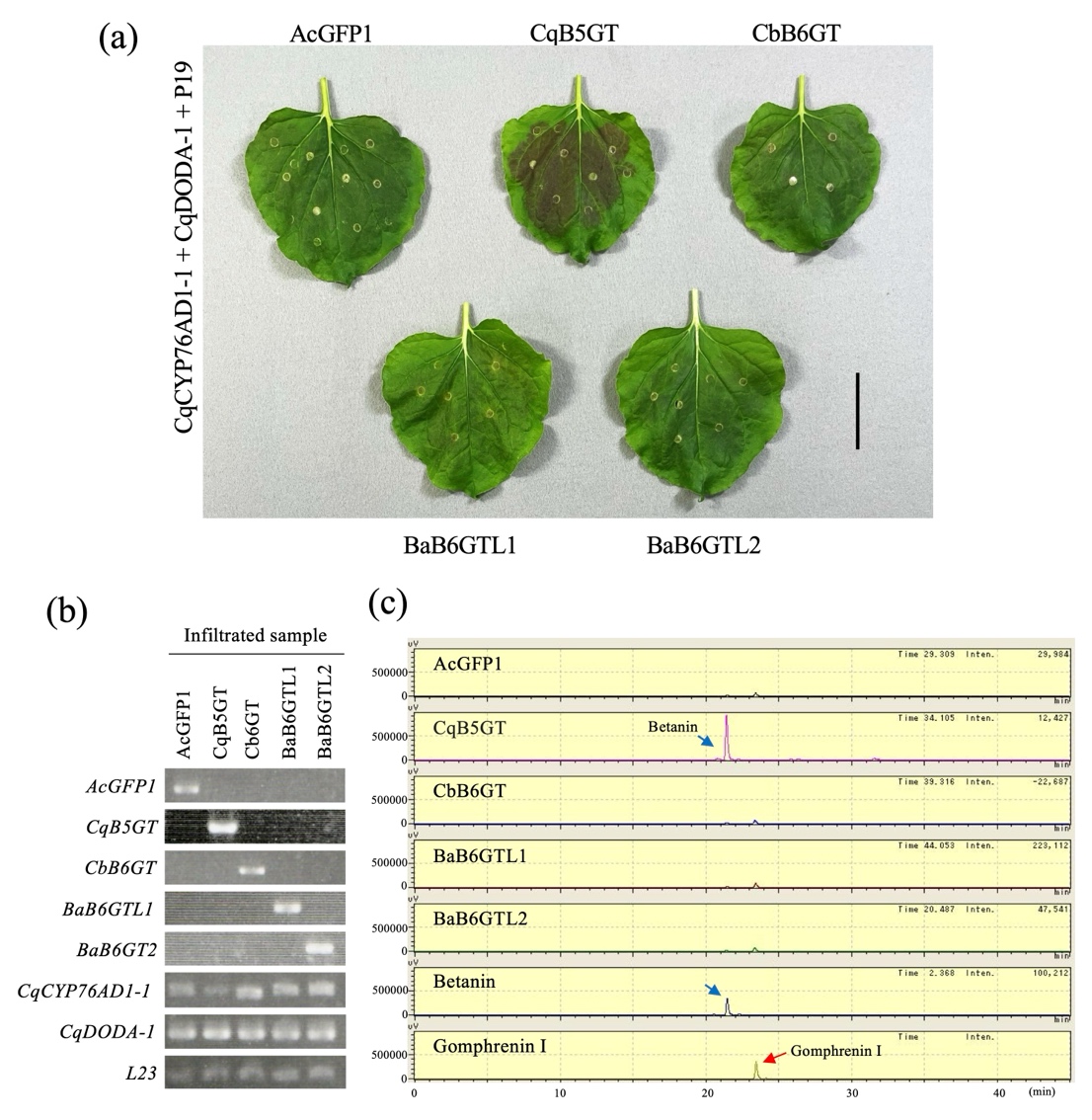

**Figure S5. Identification and functional analysis** **of B6GT family genes. (a)** Recombinant expression of B6GT family genes in *Nicotiana benthamiana* leaves. *CbB6GT*, *BaB6GTL1*, and *BaB6GTL2* indicate co-infiltration of transgenic *Agrobacterium* strains harboring plasmids containing each B6GT gene (*CbB6GT*, *BaB6GTL1*, or *BaB6GTL2*), together with *CqCYP76AD1-1*, *CqDODA-1*, and *P19*. *CqB5GT* was used as a positive control for betanin production, confirming that the betalain biosynthetic system was functional. *AcGFP1* was used as a negative control. An additional control lacking AcGFP1 (CqCYP76AD1-1 + CqDODA-1 + P19) is shown in Figure S6 to evaluate the effect of AcGFP1 on pigmentation. Bar, 4 cm. **(b)** RT-PCR analysis of transgene expression in infiltrated *N. benthamiana* leaves. *L23* served as the internal control. **(c)** HPLC chromatograms of extracts from infiltrated *N. benthamiana* leaves. The red and blue arrows indicate the peaks corresponding to gomphrenin I and betanin, respectively. The horizontal axis represents the retention time (min), and the vertical axis represents the signal intensity (µV).

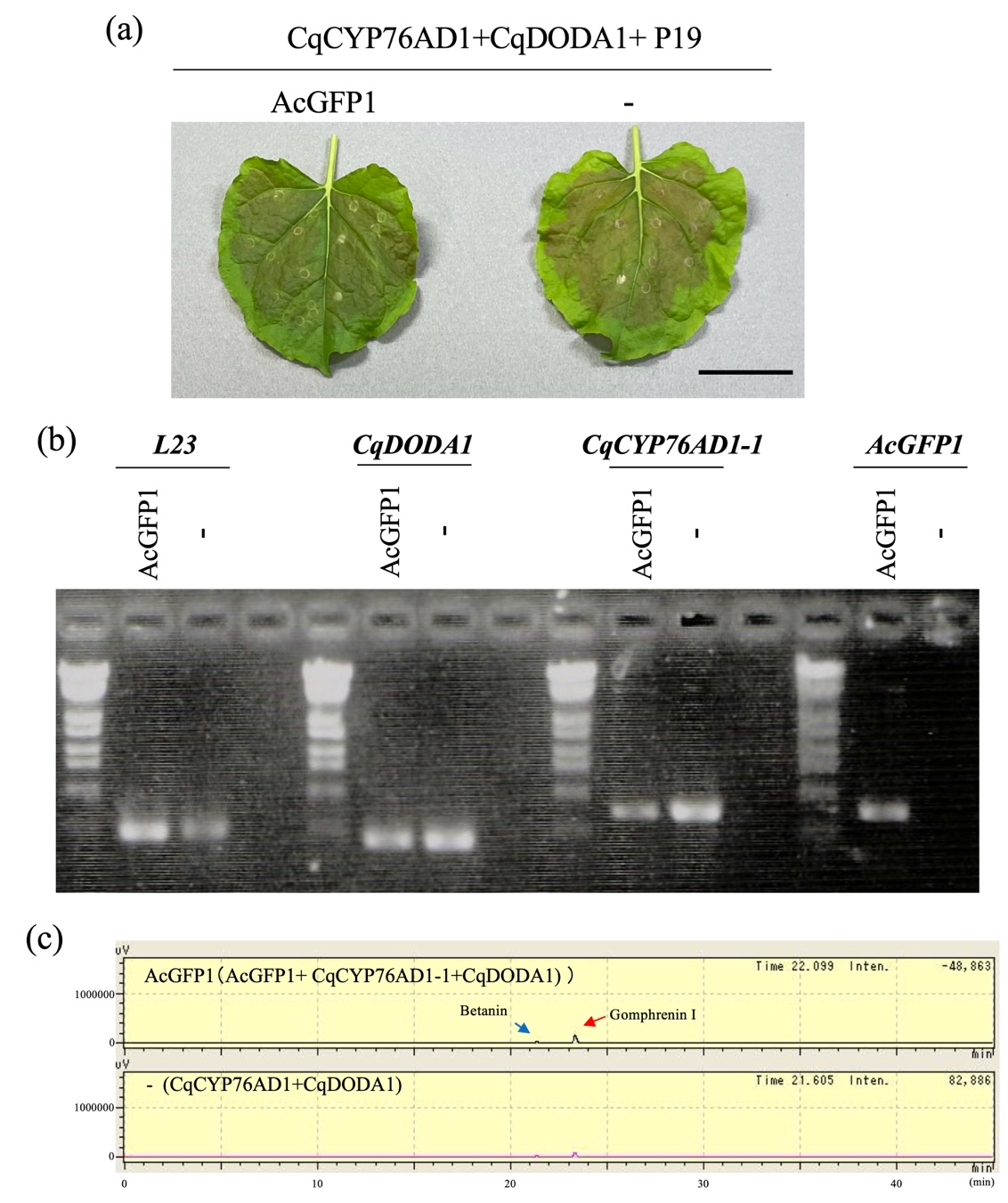

**Figure S6. Evaluation of UGT-free control conditions in *Nicotiana benthamiana.* (a)** Transient expression of the control gene set in *N. benthamiana* leaves. *Agrobacterium* strains harboring AcGFP1 were co-infiltrated with *CqCYP76AD1-1*, *CqDODA-1*, and *P19*. The minus sign (−) indicates samples infiltrated with *CqCYP76AD1-1*, *CqDODA-1*, and *P19* only (i.e., without AcGFP1). Bar, 4 cm. **(b)** RT-PCR analysis of transgene expression in infiltrated leaves. L23 was used as an internal control. **(c)** HPLC chromatograms of extracts from infiltrated leaves. Blue and red arrows indicate peaks corresponding to betanin and gomphrenin I, respectively. The horizontal axis represents retention time (min), and the vertical axis represents signal intensity (µV). Only trace-level peaks were detected under both conditions.

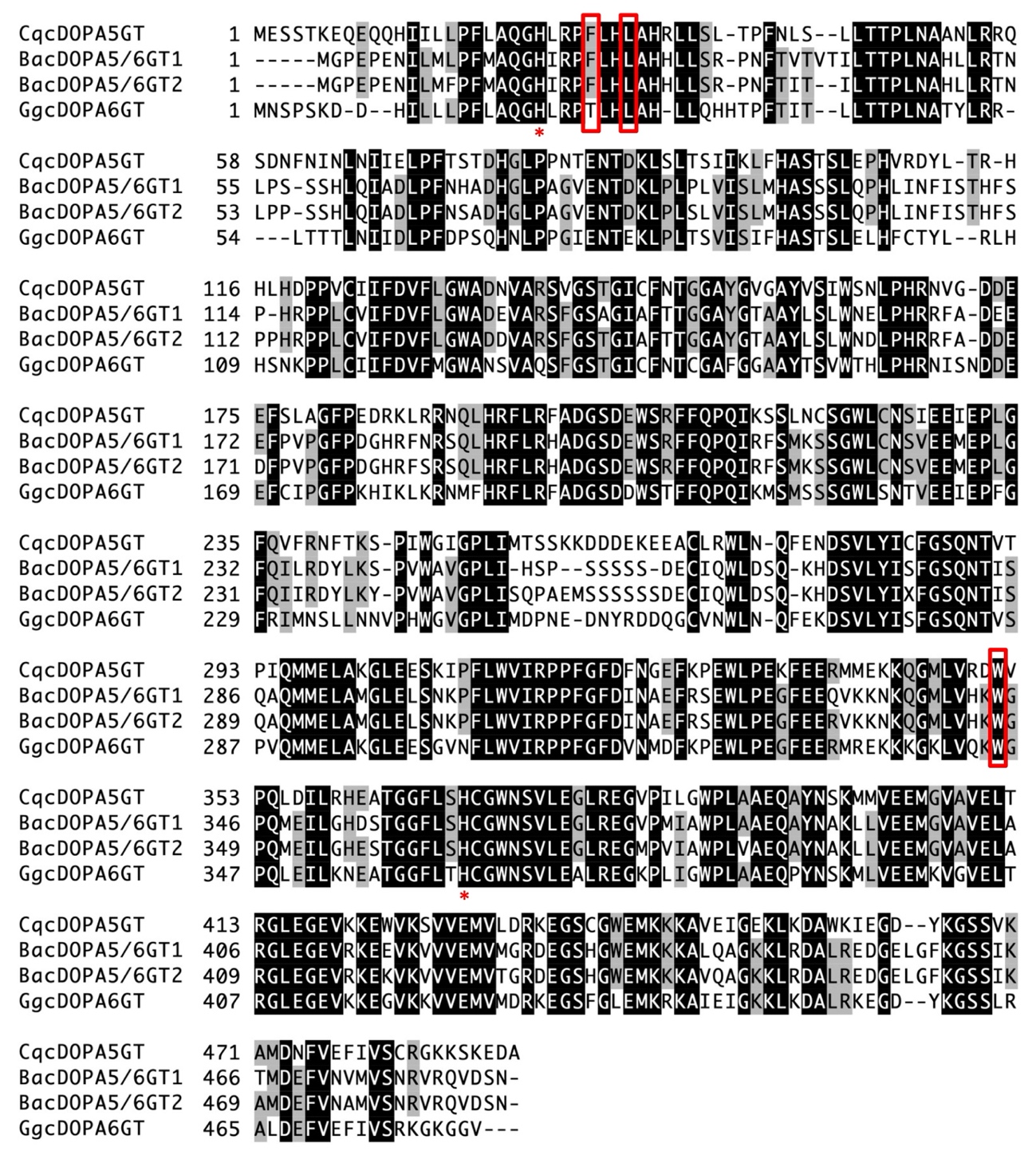

**Figure S7. Alignment of deduced amino acid sequences of the cDOPA-glucosyltransferase family.** The cDOPA-GTs used for the alignment were derived from C. quinoa, B. alba, and G. globosa. Asterisks indicate amino acid residues predicted to be part of the catalytic site. Red boxes denote amino acid residues predicted by 3D simulation analysis to influence enzyme activity, although they are not located within the catalytic site.

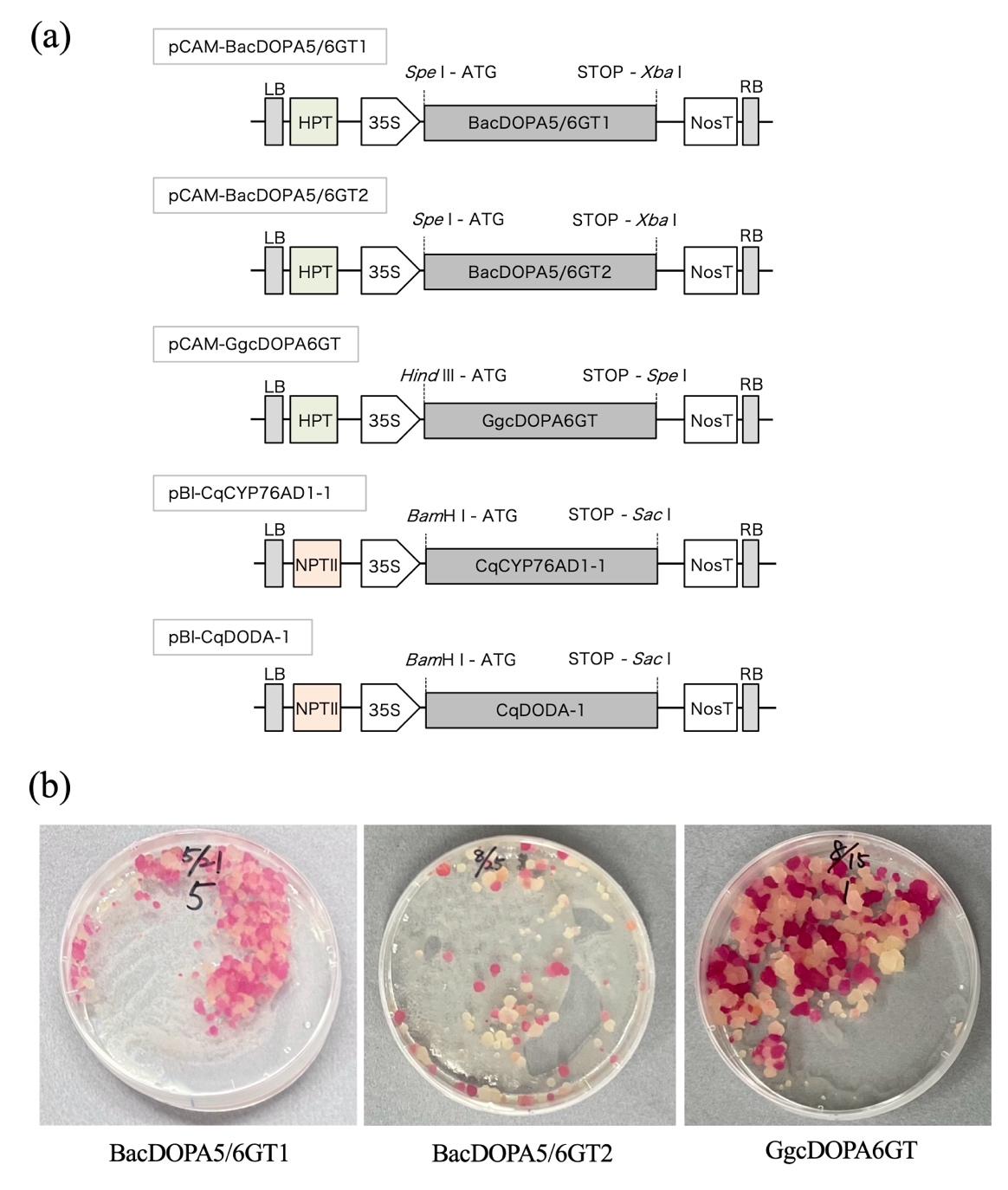

**Figure S8. Generation of betalain-producing BY-2 cell lines. (a)** Schematic representations of the plant expression vector. BacDOPA5/6GT1, *BacDOPA5/6GT1* CDS; BacDOPA5/6GT2, *BacDOPA5/6GT2* CDS; GgcDOPA6GT, *GgcDOPA6GT* CDS; CqCYP76AD1-1, *CqCYP76AD1-1* CDS; CqDODA-1, *CqDODA-1* CDS; 35S, CaMV 35S promoter; NosT, *nopaline synthase* terminator; 35S-T, 35S terminator; RB, right border; LB, left border; HPT, *hygromycin phosphotransferase* expression cassette; NPTII, *neomycin phosphotransferase II* expression cassette; bar*, bar* gene expression cassette; ATG, start codon; STOP, stop codon. **(b)** Photographs of betalain-producing cell lines taken **three weeks after infection**. The left, middle, and right panels show *BacDOPA5/6GT1-*, *BacDOPA5/6GT2*-, and *GgcDOPA6GT-*expressing lines, respectively. **Red colonies indicate independent transgenic lines.**

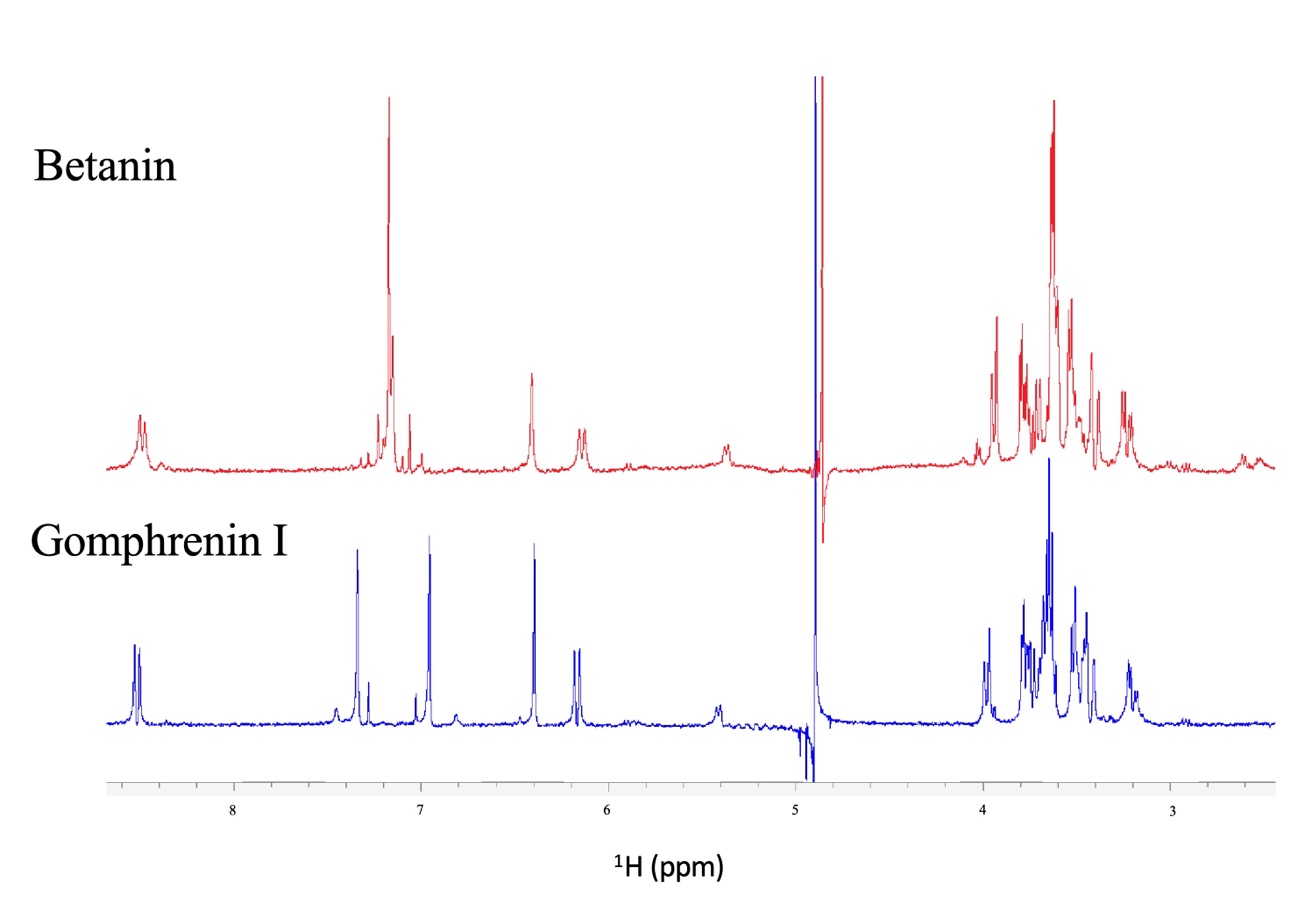

**Figure S9. ^1^H-NMR spectra of gomphrenin I and betanin.** The upper and lower spectra correspond to betanin and gomphrenin I, respectively.

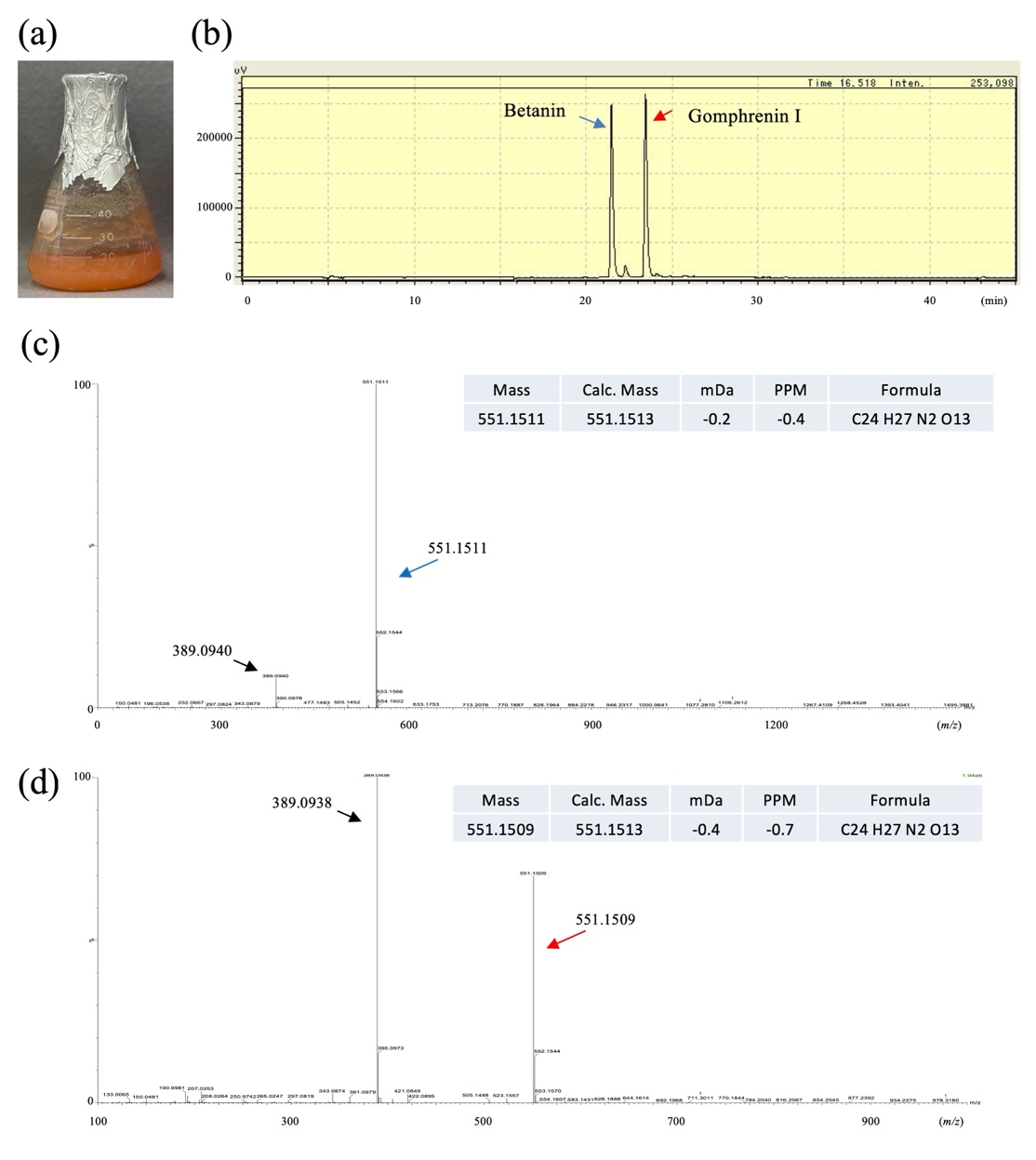

**Figure S10. Pigment analysis of the betanidin-producing BY-2 cell line. (a)** Photograph of the Betanidin line, corresponding to BY-2 cells transformed with CqCYP76AD1-1 and CqDODA-1. Cells were cultured to a high-density condition prior to pigment extraction. **(b)** HPLC chromatogram of pigment extracts from the Betanidin line. Pigments were extracted from a large volume of cells (~1 L culture) to obtain sufficient material for analysis. Blue and red arrows indicate betanin and gomphrenin I, respectively. The horizontal axis represents retention time (min), and the vertical axis represents signal intensity (µV). **(c, d)** Mass spectra corresponding to the HPLC peaks indicated in (b). Upper and lower panels show spectra from the 21–22 min peak (blue arrow) and the 23–24 min peak (red arrow), respectively. Blue and red peaks correspond to betanin and gomphrenin I, respectively. The horizontal axis indicates mass-to-charge ratio (*m/z*), and the vertical axis indicates relative abundance. Arrows indicate characteristic betanidin-derived fragment ions.

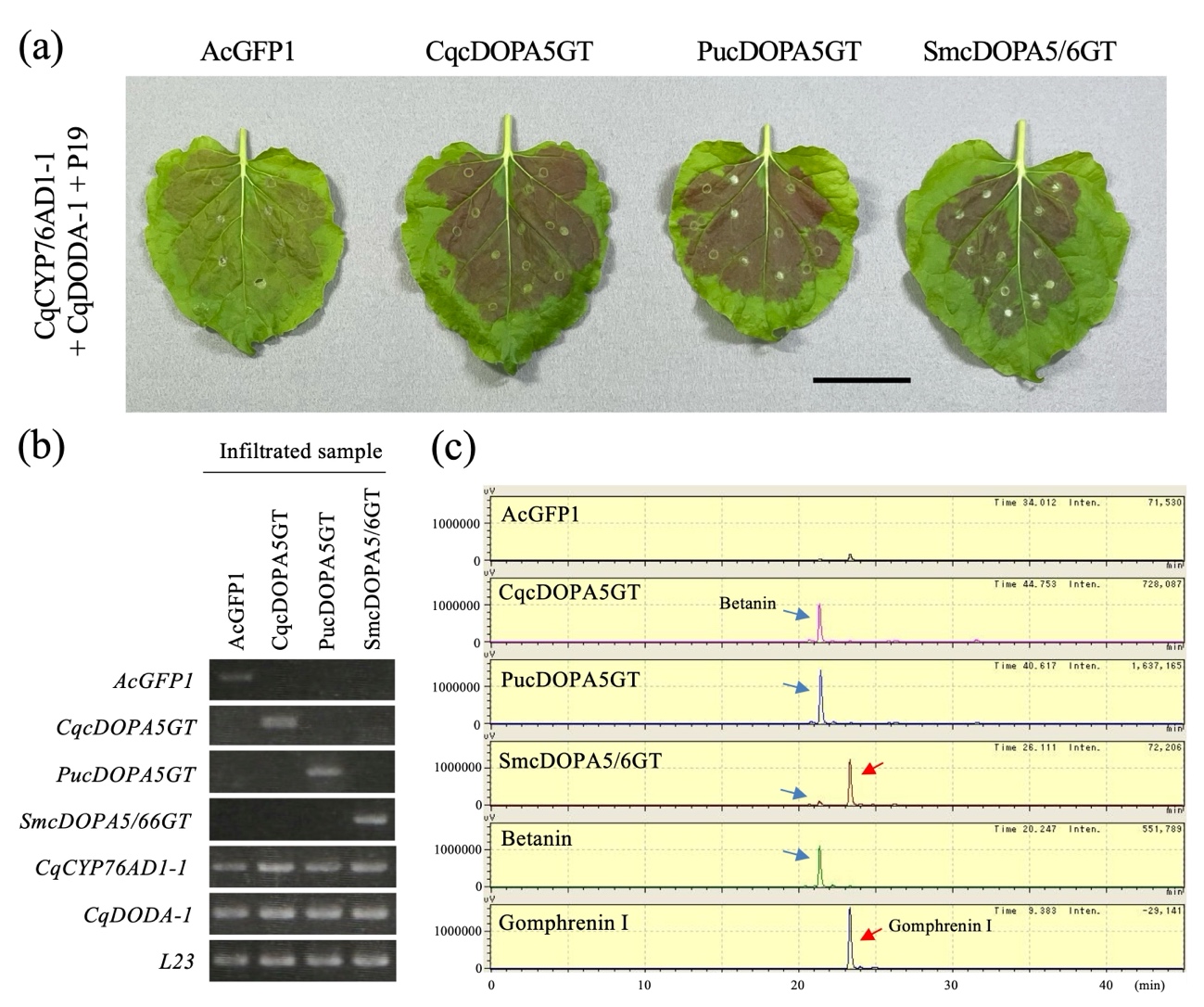

**Figure S11. Functional characterization of cDOPA6GT-related enzymes from other plant species.** **(a)** Recombinant expression of cDOPA5GT and cDOPA5/6GT-related enzymes in Nicotiana benthamiana leaves. Transgenic Agrobacterium strains harboring plasmids carrying *PucDOPA5GT*, *SmcDOPA5/6GT*, or *CqcDOPA5GT* were co-infiltrated with *CqCYP76AD1-1*, *CqDODA-1*, and *P19*. *AcGFP1* was used as a negative control. Bar, 4 cm. **(b)** RT-PCR analysis of transgene expression in infiltrated *N. benthamiana* leaves. *L23* served as the internal control. **(c)** HPLC chromatograms of extracts from infiltrated N. benthamiana leaves. Betanin and gomphrenin I standards are shown for comparison. The red and blue arrows indicate the peaks corresponding to gomphrenin I and betanin, respectively. The horizontal axis represents retention time (min), and the vertical axis represents signal intensity (µV).

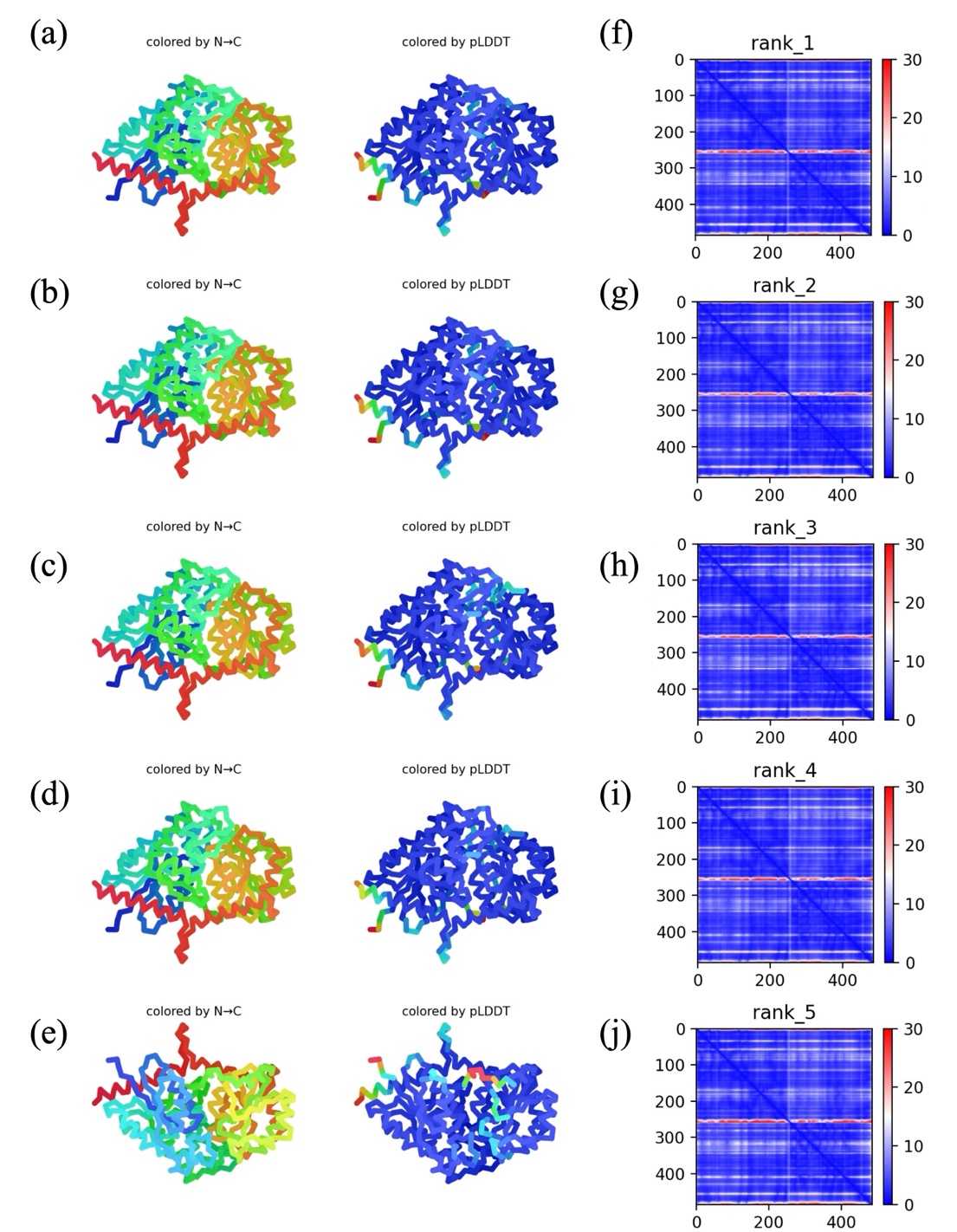

**Figure S12. AlphaFold structural models of BacDOPA5/6GT1. (a–e)** Predicted structures of BacDOPA5/6GT1 corresponding to rank 1–5 models. Structures are shown and colored by residue position (N-terminus to C-terminus) and by prediction confidence (pLDDT), where blue indicates high confidence and red indicates low confidence. **(f–j)** Predicted Aligned Error (PAE) plots corresponding to each model (rank 1–5). The PAE values indicate the expected positional error between residue pairs, with lower values (blue) representing higher confidence in the relative positioning of residues. All models were generated using AlphaFold. pTM scores are indicated in the Supplementary Data. ipTM scores are not applicable, as the models correspond to single-chain proteins.

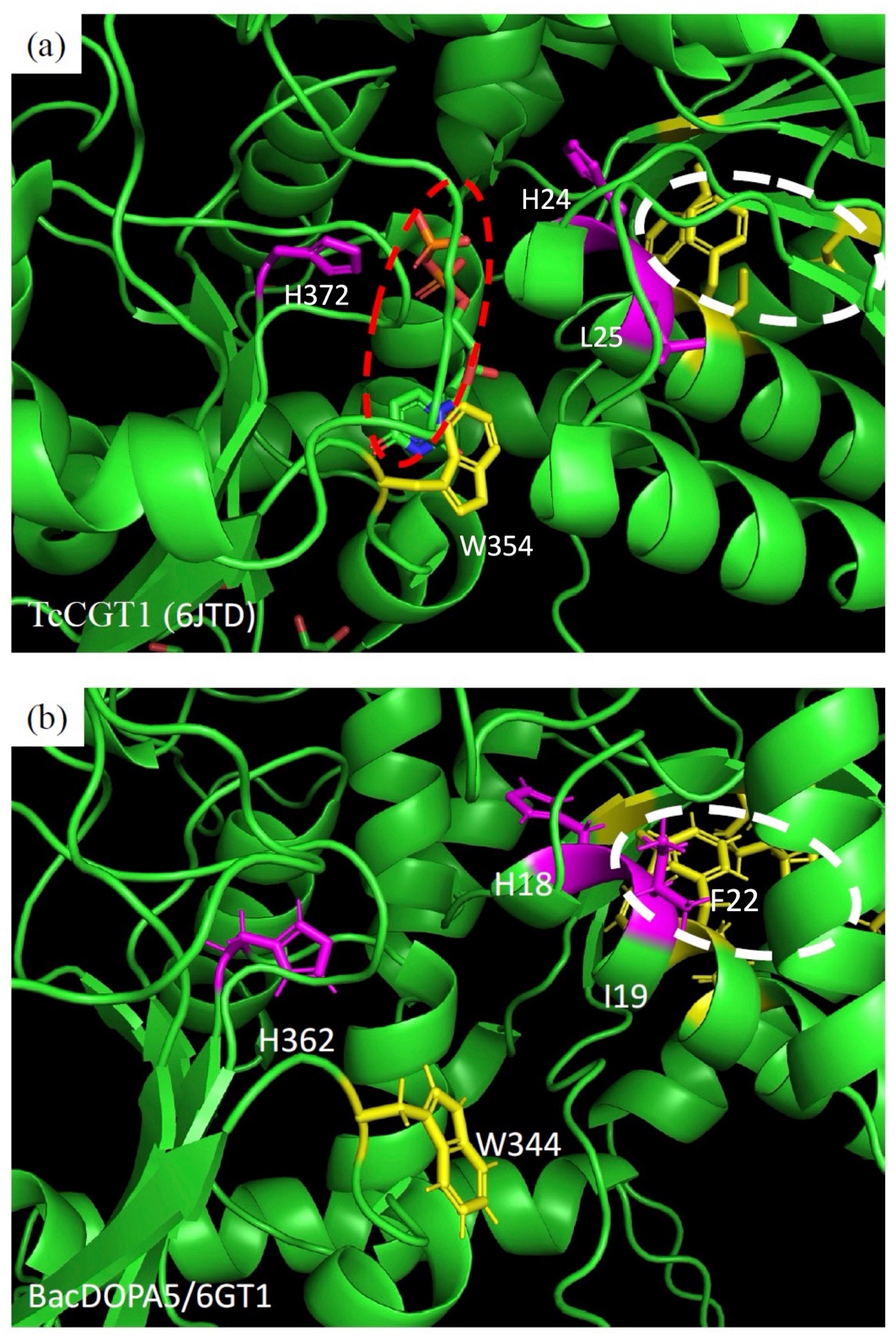

**Figure S13. Comparison of the crystal and simulated catalytic-site** **structures in glycosyltransferases. (a, b)** Three-dimensional structures of the catalytic sites of *Trollius chinensis* C-glycosyltransferase 1 (*TcCGT1*, PDB ID: 6JTD) **(a)** and *BacDOPA5/6GT1* **(b)**. Red dashed circles indicate the substrate molecules, and white dashed circles denote clusters of hydrophobic residues formed by α-helices and β-sheets.

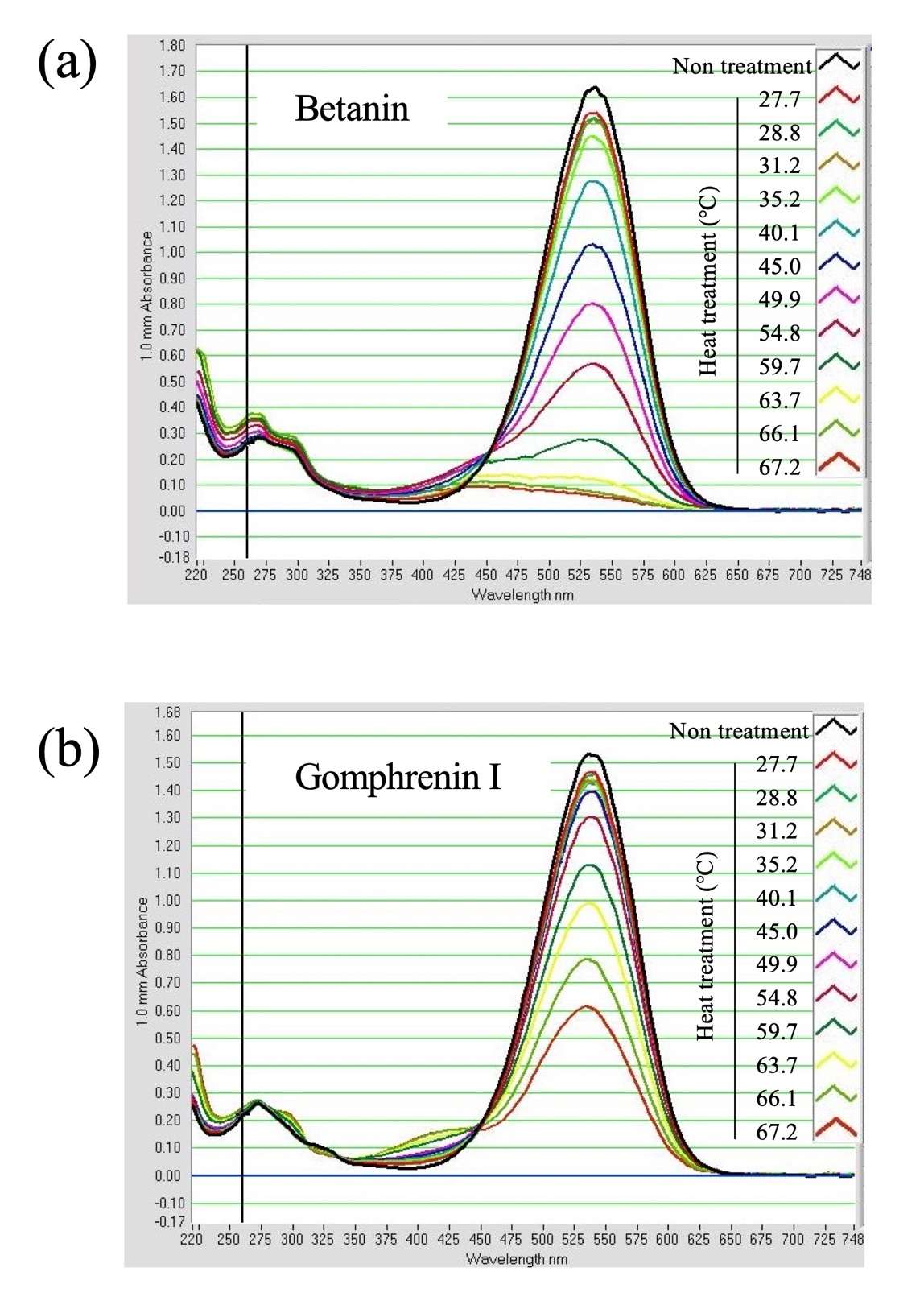

**Figure S14. Heat stability of gomphrenin I and betanin.** Absorption spectra of heat-treated betanin (a) and gomphrenin I (b). The vertical axis shows absorbance, and the horizontal axis indicates wavelength (nm).

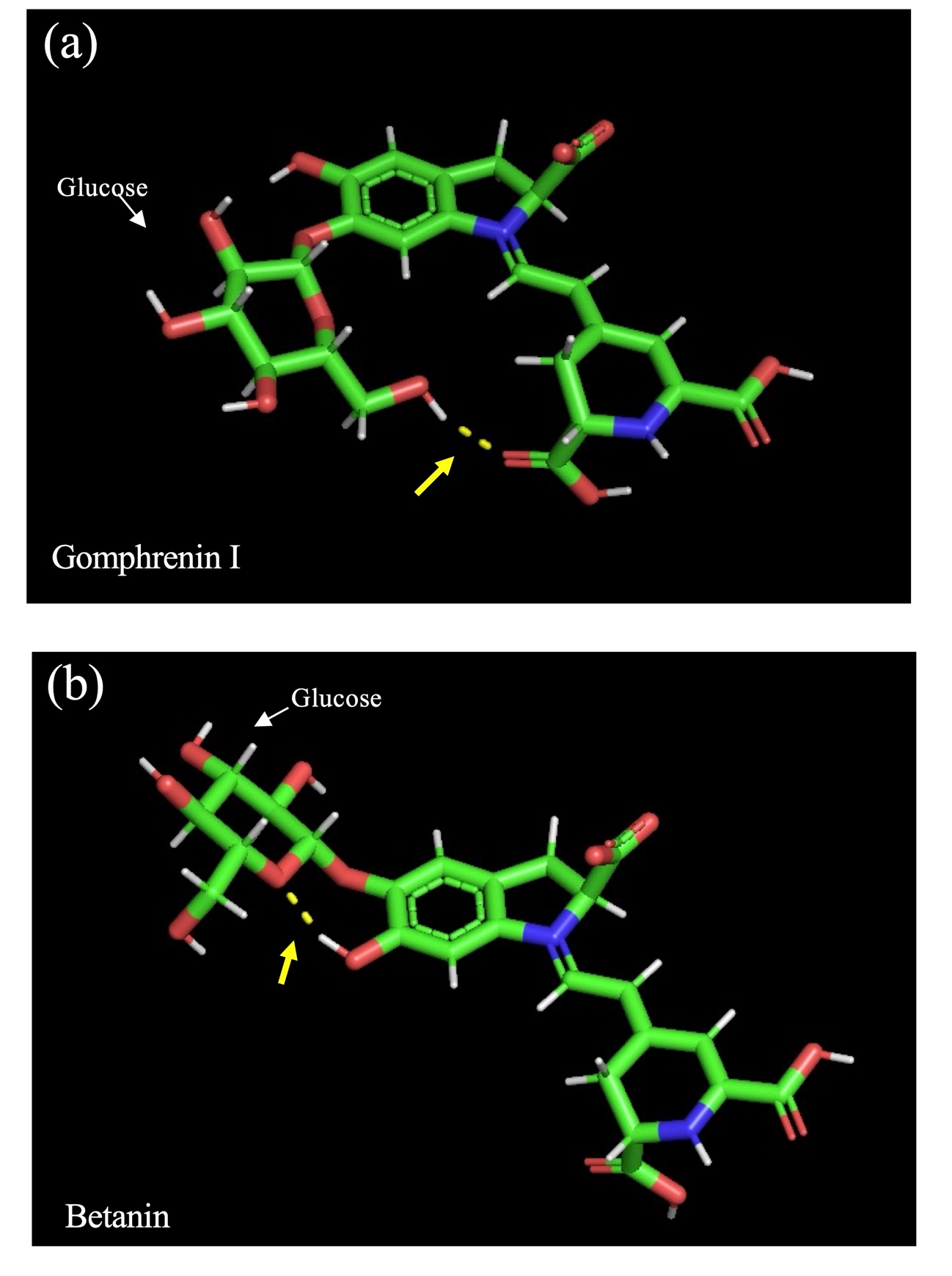

**Figure S15. Molecular structures of gomphrenin I and betanin generated by molecular orbital calculations.** (a) Gomphrenin I; (b) betanin. Yellow dashed lines indicate a possible intramolecular hydrogen bond predicted in gomphrenin I.

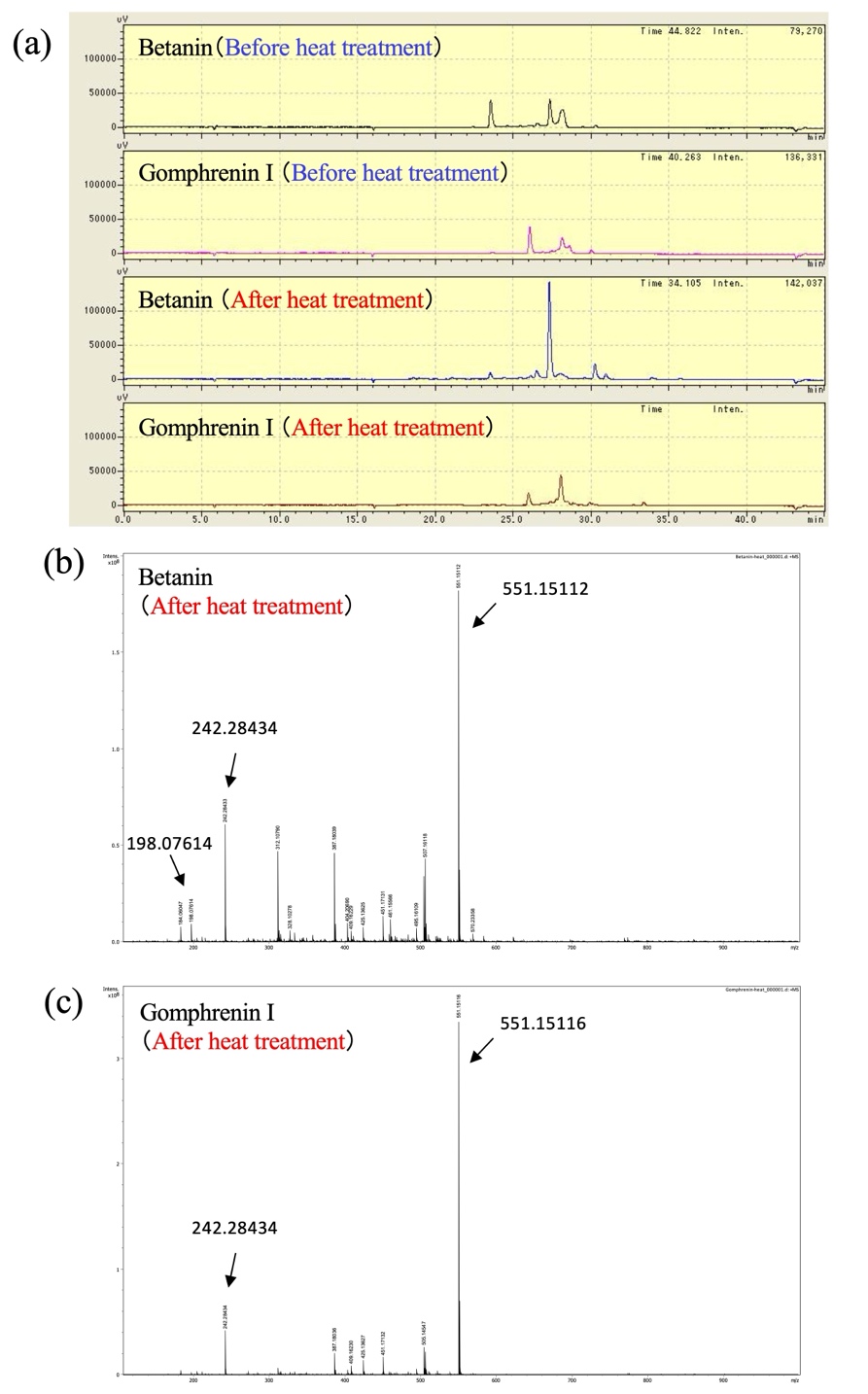

**Figure S16. HPLC and MS spectrum of betanin and gomphrenin I before and after heat treatment. (a)** HPLC chromatograms of betanin and gomphrenin I before and after heat treatment. Samples were subjected to heat treatment (65°C, 3 h). HPLC analysis was performed under standard conditions for betalain detection, with monitoring at 408 nm. **(b)** MS spectrum of betanin after heat treatment. The major ion at m/z 551.15112 corresponds to intact betanin. **(c)** MS spectrum of gomphrenin I after heat treatment. The major ion at m/z 551.15116 corresponds to intact gomphrenin I. Notably, no prominent signal corresponding to betalamic acid (expected m/z 211) was detected under these conditions.

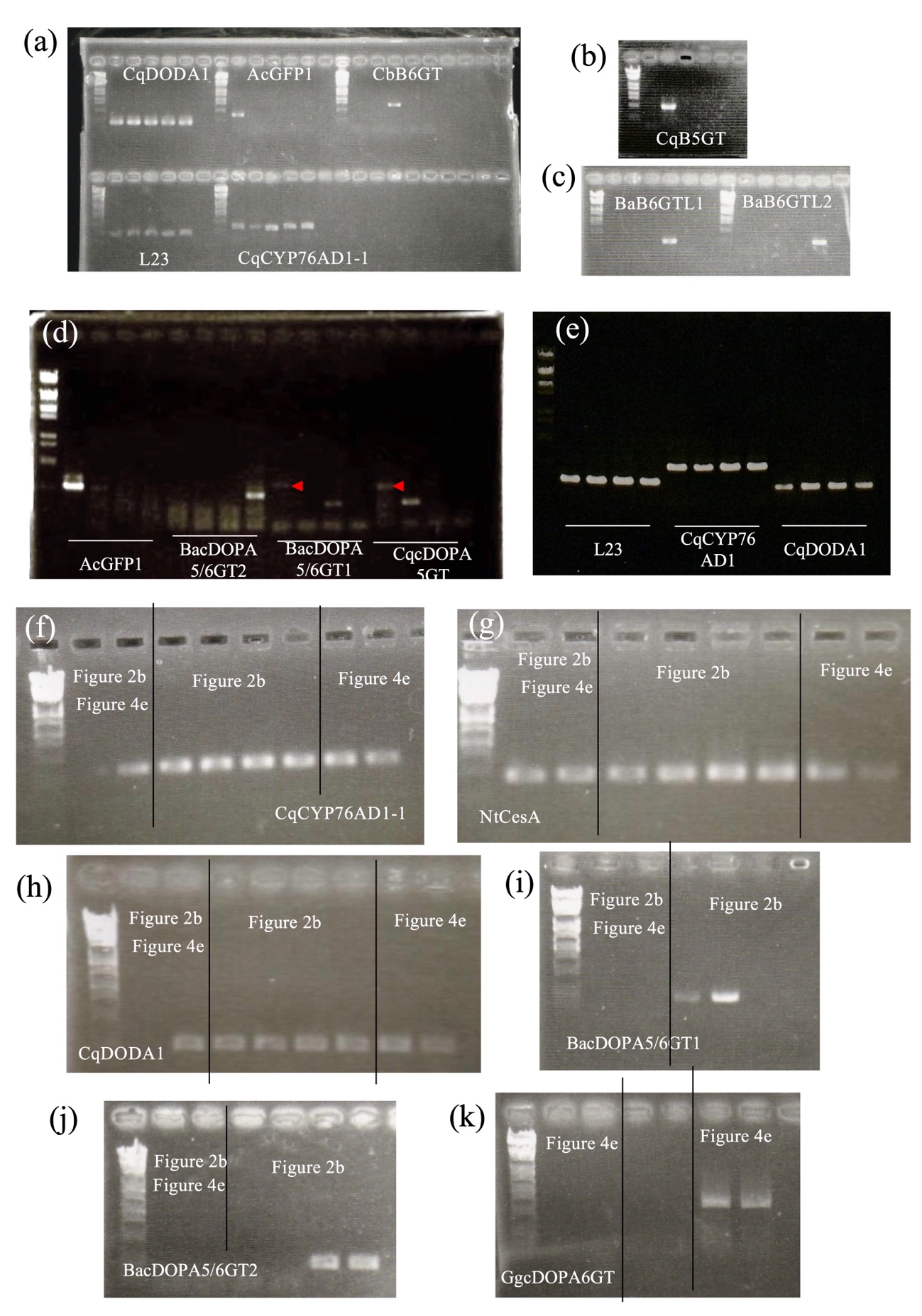

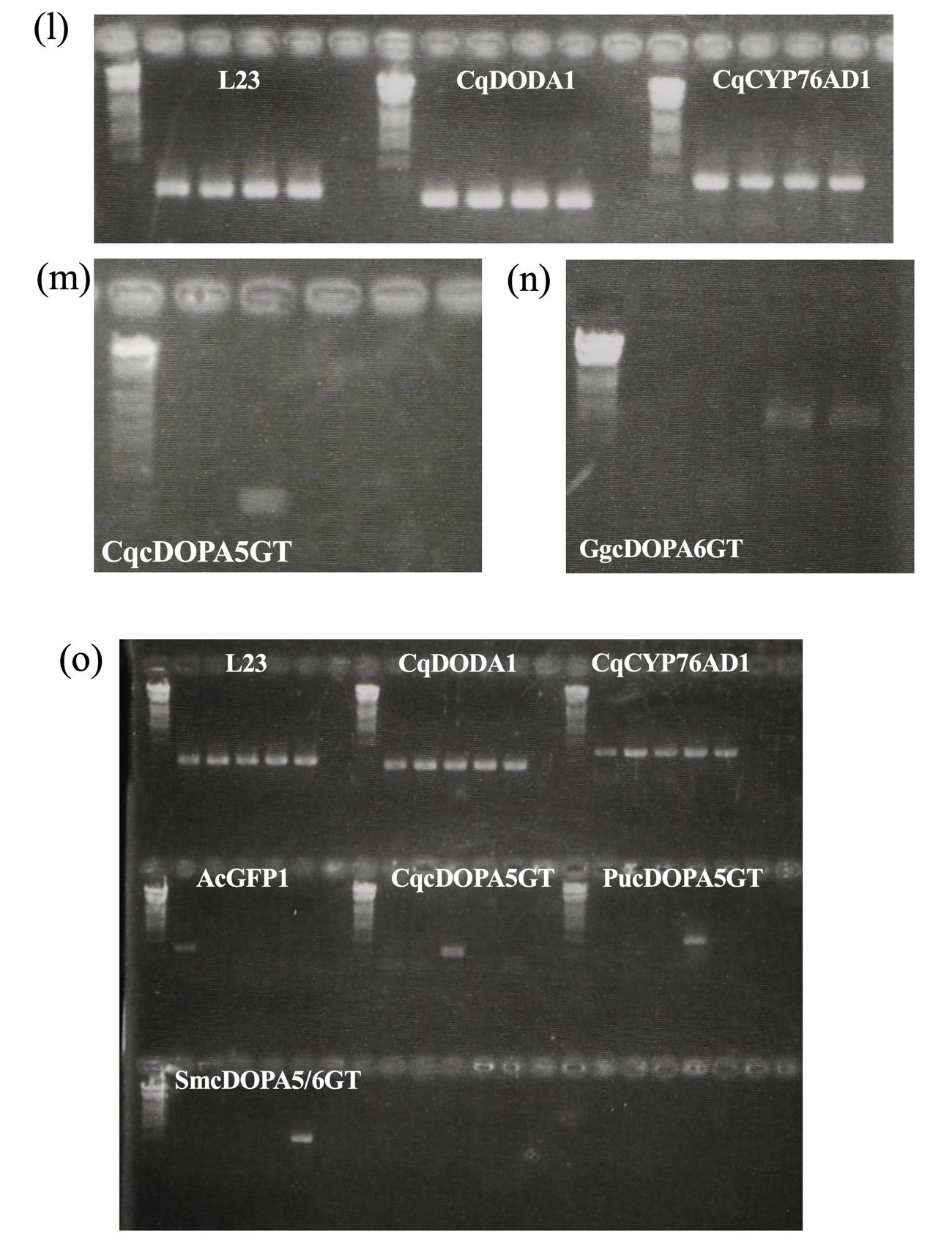

**Figure S17. Uncropped gel images of RT-PCR.** RT-PCR images corresponding to Figure S5 **(a–c)**, Figure 2b **(d, e)**, Figure 3b, Figure 5e **(f–k)**, Figure 5b **(l–n)**, and Figure S11 **(o)** are shown. DNA size markers (λ DNA digested with EcoT14I) are included. The red arrowhead indicates a non-specific PCR amplification band.
